# Allergen-induced airway matrix remodelling in mice can be prevented or reversed by targeting chitinase-like proteins

**DOI:** 10.1101/2023.08.18.553857

**Authors:** James E Parkinson, Antony Adamson, Andrew S MacDonald, Judith E Allen, Tara E Sutherland

**Affiliations:** Lydia Becker Institute of Immunology and Inflammation, School of Biological Sciences, Faculty of Biology, Medicine and Health, University of Manchester, Manchester, M139PT, UK; Wellcome Centre for Cell Matrix Research, School of Biological Sciences, Faculty of Biology, Medicine and Health, University of Manchester, Manchester, M139PT, UK; Genome Editing Unit, Faculty of Biology, Medicine and Health, University of Manchester, Manchester, M139PT, UK; Institute of Medical Sciences, School of Medicine, Medical Sciences and Dentistry, University of Aberdeen, Aberdeen AB252ZD, UK

## Abstract

Chitinase-like proteins (CLPs) are biomarkers of inflammation and airway remodelling in asthma, yet their direct contribution towards disease pathogenesis is unknown. Using a mouse model of allergen-induced type 2/type 17 airway inflammation we sought to directly investigate the role of the murine CLPs Ym1 and Ym2 during chronic lung pathology. Data demonstrated distinct chronic inflammatory roles for Ym2, IL-13, and IL-17a signalling pathways. Notably, only CLPs were key for initiating the pathogenic accumulation and re-organisation of the pulmonary extracellular matrix (ECM) environment. Furthermore, inhibition of CLPs after chronic pathology developed, reversed airway remodelling independently of chronic inflammation. These studies disentangle chronic IL-13 and IL-17a signalling from the development of allergic airway remodelling and instead highlight a central role for CLPs, which provides new avenues to therapeutically target aberrant ECM accumulation.

## Introduction

The relationship between our immune system and the extracellular matrix (ECM) plays a key role in maintaining tissue homeostasis. The different molecules of the ECM, including fibrillar proteins, glycosaminoglycans and proteoglycans, provide a vital physical and biological support for cells and tissues in health^1^. However, changes to the organisation and composition of the lung ECM, often facilitated by dysregulation of our immune system, occurs during many respiratory disorders^2, 3^. In asthma there is compelling correlative evidence that changes or remodelling of the ECM, combined with changes to the airway epithelium, mucus hypersecretion and thickening of the airway smooth muscle layer have a detrimental impact on lung function^4–9^ and contribute to irreversible airflow obstruction^10, 11^. Whilst the complex, dynamic and heterogeneous processes of airway remodelling in asthma, including changes to the ECM, has traditionally been attributed to aberrant and persistent inflammation (reviewed in^12^), many questions remain over which specific immune cells and mediators can influence discrete features of lung remodelling in asthma.

Th2 immunity and tissue remodelling processes coincide in many conditions, and type 2 immune responses are known to regulate wound repair, including reorganisation of the ECM^13^. Whilst responses in allergic asthma can be heterogenous, they are generally considered synonymous with Th2 activation and eosinophilic inflammation^14, 15^. Therefore, not surprisingly the Th2 response in asthma is often regarded as causal of airway remodelling^16^. As such, targeting type 2 inflammatory pathways has been an attractive target for asthma therapy and commonly supported by studies in sub-chronic Th2-skewed mouse models where inhibition of type 2 signalling can prevent airway inflammation and pathology^17–22^. Nonetheless, recent systematic reviews suggest limited efficacy of modulating the Th2 response to ameliorating asthma symptoms^23, 24^. Alongside classical Th2 high asthma, asthma can also be classified as severe IL-17A, neutrophilic dominated disease characterised by airway remodelling^25^ and insensitivity to the traditionally potent anti-inflammatory steroid treatments^26^. However, targeting IL-17RA in asthma has also proved disappointing^27^. Overall, these findings have somewhat challenged the historical view that individual immune responses are causative of airway remodelling and asthma pathogenesis. Rather, more recent views suggest that remodelling may not be a result of specific immune responses, but rather occur in parallel to or even independently of inflammation^28–30^. Whilst there is no doubt that inflammatory cells and cytokines may contribute to individual airway remodelling outcomes, targeting these immune pathways alone is likely insufficient to modify the disease.

A genome-wide association study identified *CHI3L1*, the gene that encodes for chitinase-like protein YKL40, as a susceptibility gene for asthma^31^, with levels of YKL40 in the serum and tissue correlating with asthma severity, degree of remodelling and poor disease control^32–34^. Despite diversity amongst CLP members across species, all CLPs are widely associated with inflammation, host damage and tissue repair^35^. Mice express three main CLPs including Brp39, Ym1, and Ym2. Whilst Brp39 has the highest homology to human YKL40, it is Ym1 that shows more similarity in terms of both cellular expression and function^36^. Furthermore, we recently described distinct expression patterns between Ym1 and Ym2 during allergic airway pathology suggesting divergent function between these molecules despite high protein sequence similarity^37^. While research to date has largely focussed on the ability of CLPs to regulate immune responses, CLPs can also influence the ECM through direct interactions with ECM components^38–40^. We have also previously shown that direct administration of Ym1 to mice, can repair tissue damage caused by the lung migrating nematode *Nippostronglyus brasiliensis*, an effect independent of type 2 immunity^41^. Together with studies showing that YKL40 levels correlate with degree of airway wall thickening in people with asthma^34, 42^, these murine studies highlight the potential of CLPs to influence the ECM and to facilitate tissue repair and remodelling as a part of host-defence or disease pathology.

In mice, features of human disease can be modelled using chronic exposure to allergen cocktail DRA (House **D** ust Mite, **R**agweed, **A** *spergillus fumigatus*). These features include eosinophil/neutrophil inflammation, steroid insensitivity, airway remodelling and persistent pathology^37, 43, 44^. In this study we use the DRA model to not only characterise the roles of IL-17a and IL-13 in chronic allergic pathology, but also investigate whether CLPs contribute to airway remodelling responses. We show that remodelling responses, including changes to the levels and diversity of ECM components, can occur independently of chronic allergic type 2 eosinophil-driven or IL-17a neutrophil-driven inflammation. Instead, mice administered chronic DRA allergens only developed a fully cross-linked ECM pathogenic matrix in the presence of secreted Ym2. Remarkably, therapeutic inhibition of Ym1 in mice with stable allergic airway disease reversed not only the pathogenic ECM remodelling response, but also the increase in airway muscle mass. Together, these findings provide a mechanistic understanding of how Ym1 and Ym2 contribute to the development of inflammation-independent allergic remodelling and support the notion of targeting CLPs in lung diseases that feature aberrant ECM accumulation.

## Results

### IL-17a is required for allergen-induced eosinophilia and neutrophilia but dispensable for airway remodelling

We previously demonstrated a different in immune signatures between C57BL/6 and BALB/c administered DRA allergens^37^. Despite this, allergic airway remodelling developed in both strains of mice, suggesting a potential disconnect between inflammation and remodelling. IL-17a levels are significantly increased in the DRA model^37^ and IL-17a is associated with severe asthma phenotypes in people^26^. Therefore, we specifically evaluated the role of IL-17a responses during DRA-induced allergic pathology by using *Il17a*^Cre^Rosa26^eYFP^ mice^45^. We validated that these mice expressed Cre in place of the *Il17a* gene resulting in *Il17a* gene deficiency when homozygous. Cre excision of a stop cassette resulted in YFP expression from the Rosa locus and generated a marker of cells in which *Il17a* would ordinarily be expressed in the DRA model (**Fig 1a-c**). As IL-17 responses are known to contribute to neutrophilic inflammation, we observed an expected increase in neutrophil numbers in the BAL in wild-type but not IL-17a-deficient mice administered DRA allergens (**Fig 1d**). However, IL-17a was also required for an established eosinophilia (**Fig 1d**) and type 2 inflammatory responses (**Fig 1e**). The link between IL-17a signalling and development of type 2 responses was entirely consistent with our previous findings during helminth infection whereby IL-17a suppression of IFNɣ is needed for the induction of pulmonary type 2 immunity^46, 47^. Despite reduced type 2 immune responses, equivalent goblet cell hyperplasia was observed in both wild-type and IL-17a-deficient allergic mice (**Fig 1f**). Additionally, preventing chronic inflammation in allergic IL-17a-deficient animals was not enough to limit the accumulation of airway collagen (**Fig 1g**). Furthermore, increased expression of ECM molecules collagen I, III or glycosaminoglycan hyaluronan (HA) around the airways following DRA administration was evident in both wild-type and IL-17a-deficient DRA mice compared to PBS treated animals (**Fig 1h, 1i**). No change in collagen IV was observed in either wild-type or IL-17a-deficient DRA mice compared to PBS treated animals (**Fig 1h, 1i**). Increased airway smooth muscle mass and number of vimentin positive cells around the airways, another feature of extensive lung remodelling, were also not regulated by IL-17a-deficiency (**Fig 1k**). Therefore, despite being required for induction and maintenance of allergic neutrophilic and eosinophilic inflammation, IL-17a was entirely dispensable for pathogenic airway accumulation of ECM components during chronic allergic airway inflammation.

**Fig 1:**
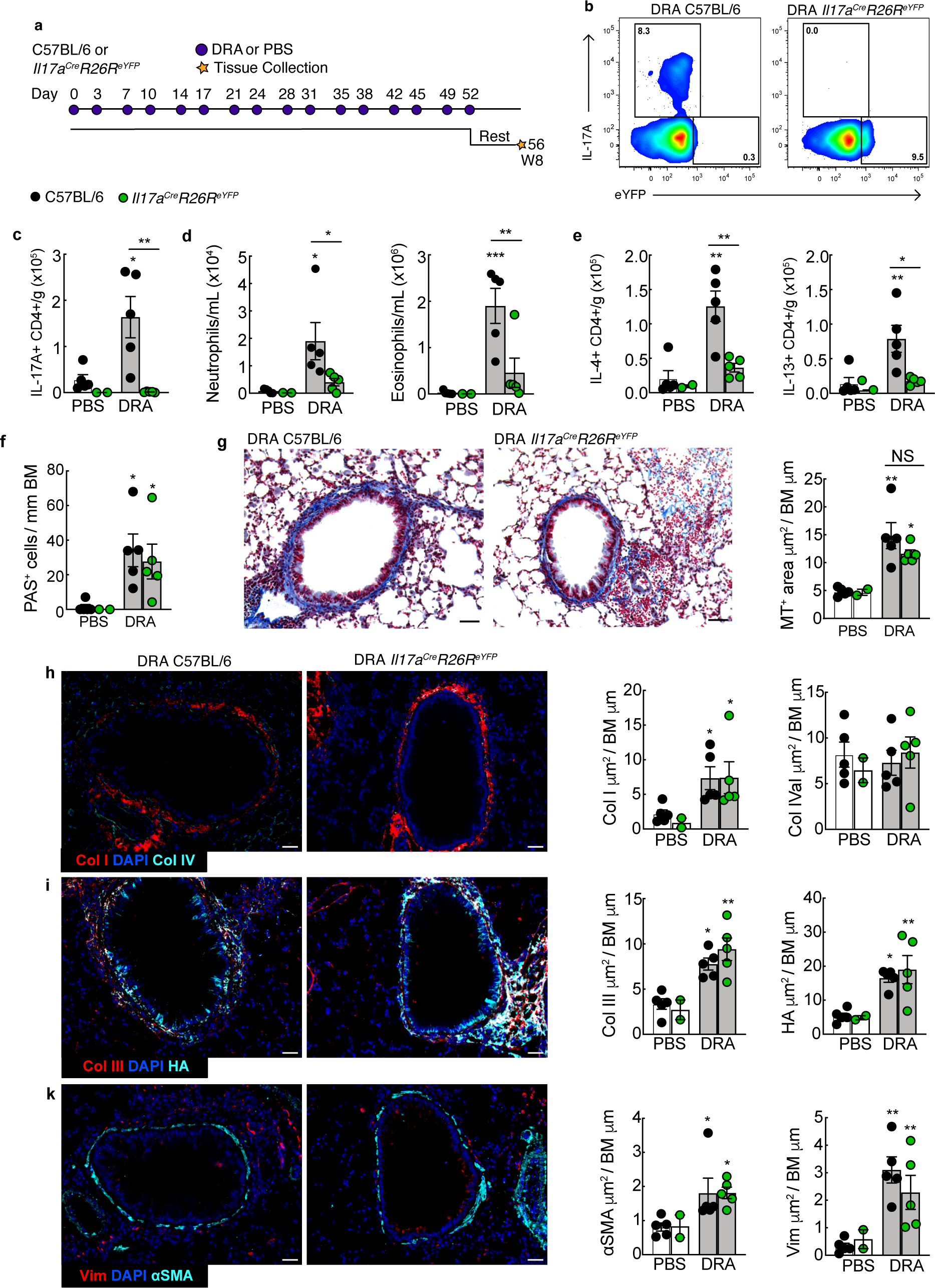
IL-17a is required for allergic inflammation but not airway ECM accumulation. **a)** DRA or PBS dosing strategy for *Il17a*^Cre^*Rosa26*^eYFP^ or C57BL/6 female mice. **b)** Representative flow cytometric identification of intracellular expression of IL-17a or eYFP signal and **c)** numbers per gram of lung of CD45^+^ TCR**β**^+^ CD4^+^ T cells from *Il17a*^Cre^*Rosa26*^eYFP^ or C57BL/6 mice administered DRA stimulated ex vivo with PMA/ionomycin. **d)** Numbers of neutrophils and eosinophils per mL of BAL in mice administered PBS or DRA according to a) and measured by flow cytometry. **e)** Numbers of IL-4^+^ or IL-13^+^ CD45^+^ TCRβ^+^ CD4^+^ T cells from *Il17a*^Cre^*Rosa26*^eYFP^ or C57BL/6 mice administered PBS or DRA stimulated ex vivo with PMA/ionomycin. Cells were analysed by flow cytometry and expressed per g of lung tissue. **f)** Lung tissue sections from mice in **a)** were stained with PAS and numbers of PAS^+^ airway epithelial cells counted per airway and normalised to length of the basement membrane. **g)** Representative histological staining of Masson’s trichrome (MT) stained tissue sections (scale bar=50μm) from mice in **a)**. MT^+^ area was analysed around the airway excluding vascular regions and normalised to basement membrane length. Representative images of lung tissue sections from DRA *Il17a*^Cre^*Rosa26*^eYFP^ or C57BL/6 mice stained with DAPI to visualise cell nuclei and antibodies or binding proteins recognising **h)** collagen I and IV, **i)** collagen III and HA binding protein (HABP) or **k)** ⍺SMA and vimentin (scale bar = 30μm). Positive-stained area around the airways were analysed for each antigen and normalised to length of basement membrane. Datapoints show individual animals with bars representing mean ± s.e.m with n=2-5 female mice per group and data are representative from two individual experiments. Data were analysed by ANOVA with Tukey’s multiple comparison test and significance level shown relative to PBS C57Bl/6 mice or between C57BL/6 and *Il17a*^Cre^*Rosa26*^eYFP^ mice as indicated on the graph. * *P*<0.05, ** *P*<0.05, *** *P*<0.05 and NS, not significant.

### IL-13 promotes activation of the airway epithelium but is dispensable for allergen-induced ECM accumulation

The type 2 cytokine IL-13 is described as a central mediator of epithelial cell function, mucus production and collagen homeostasis^48–51^. Despite a reduction of eosinophils as well as IL-4^+^ and IL-13^+^ T cells in allergic IL-17a-deficient mice (**Fig 1**), we could not exclude the possibility that IL-13 was contributing to airway pathology in these mice. Therefore, we sought to investigate the contribution of IL-13 to allergic remodelling responses using homozygous Il-13^eGFP^ reporter mice which are IL-13-deficient^52^. Chronic DRA administration (**Fig 2a**) increased IL-13-producing CD4^+^ T cells in wild-type mice or eGFP^+^ CD4^+^ T cells in Il-13 reporter mice compared to PBS (**Sup Fig 1a**). IL-13-deficiency had no impact on allergen-induced eosinophilia (**Fig 2b**) or numbers of IL-4- producing T cells (**Fig 1c**), consistent with a requirement for both IL-4 and IL-13 signalling in such type 2 responses^53^. However, airway neutrophil numbers were significantly reduced in *Il13*^−/−^ compared to *Il13*^+/+^ allergic mice, despite no change in IL-17a^+^ CD4^+^ T cells (**Fig 2b and 2c**). In line with the ability of IL-13 to remodel the airway epithelium^49, 54^, *Il13*^−/−^ DRA mice had significantly reduced numbers of PAS^+^ epithelial cells compared to *Il13*^+/+^ DRA mice (**Fig 2c**). Despite IL-13 dependent changes to the epithelium, IL-13-deficiency did not impact on DRA-mediated collagen accumulation, numbers of vimentin^+^ cells or airway smooth muscle mass (**Sup Fig 1b** & **Fig 2d**, **2e**). By contrast, airway HA levels were significantly reduced in both IL-13-heterozygote or IL-13-deficient allergic mice in this chronic lung pathology model, supporting our previous finding of IL-13 as a key regulatory of HA in acute lung disease^55^. Together, our studies point to a role for IL-13 regulation of epithelial responses and HA accumulation but not other airway remodelling features.

**Fig 2:**
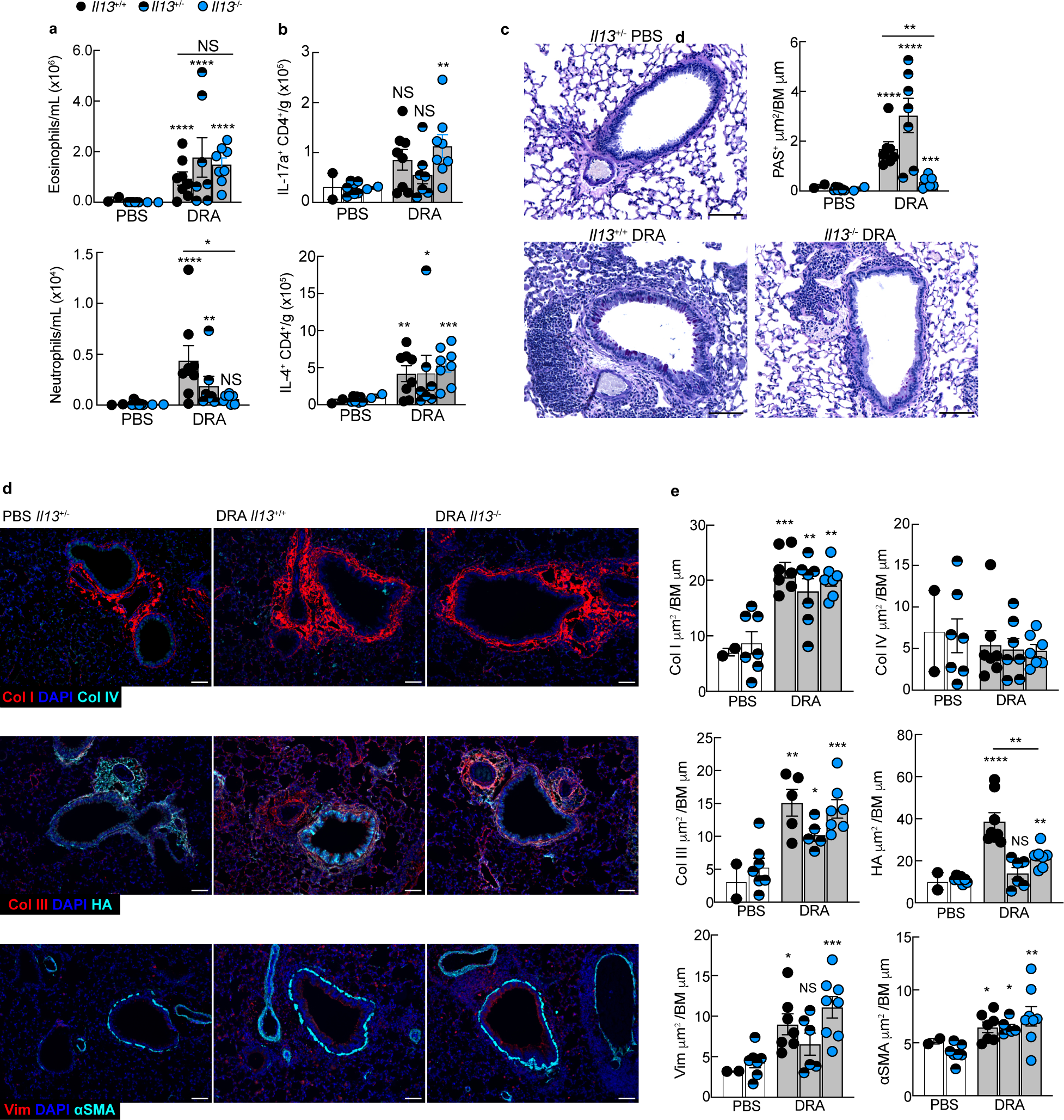
IL-13 is a driver of goblet cell hyperplasia and HA accumulation but not other remodelling features. *Il13*^eGFP^ wild-type, heterozygote or homozygote female mice were administered PBS or DRA intranasally twice a week for 8 weeks before BAL and lung tissue samples were collected 5 days after the last allergen challenge. **a)** Numbers of neutrophils and eosinophils per mL of BAL in mice as measured by flow cytometry. **b)** Numbers of IL-4^+^ or IL-17a^+^ CD45^+^ TCRβ ^+^ CD4^+^ T cells from *Il13*^eGFP^ mice administered PBS or DRA stimulated ex vivo with PMA/ionomycin. Cells were analysed by flow cytometry and expressed per g of lung. **c)** Representative histological staining of PAS^+^ airway epithelial cells in lungs from PBS or DRA treated *Il13*^+/+^ and DRA treated *Il13*^−/−^ mice (scale bar=100 μm). Graph shows the area of PAS^+^ epithelial cells normalised to basement membrane length. **d)** Representative images of lung tissue sections from PBS or DRA treated *Il13*^eGFP^ mice stained with DAPI to visualise cell nuclei and antibodies or binding proteins recognising collagen I and IV; collagen III and HA binding protein (HABP); or ⍺SMA and vimentin (scale bar = 100μm). **e)** Positive-stained area around the airways were analysed for each antigen and normalised to length of basement membrane. Datapoints show individual animals with bars representing mean ± s.e.m with n=2-7 female mice per group and data are from two combined experiments. Data were analysed by ANOVA with Tukey’s multiple comparison test and significance level shown relative to PBS *Il13*^+/−^ mice or between *Il13*^+/+^ and *Il13*^−/−^ DRA mice as indicated on the graph. \**P*<0.05, ** *P*<0.01, *** *P*<0.001, **** *P*<0.0001 and NS, not significant.

### Chit inase-like proteins are highly upregulated during chronic allergic inflammatory responses even in the absence of IL-17 or IL-13 signalling

Chitinase-like proteins are strongly associated with severe asthma and highly upregulated in animal models of allergic airway inflammation and asthma pathology^56^. Previously, we described increased expression of murine CLPs, *Chil1* (Brp-39), *Chil3* (Ym1), and *Chil4* (Ym2) following DRA administration and coinciding with ECM remodelling responses^37^. CLPs are often thought of as type 2 dependent effector molecules. However, their expression is also influenced by many other stimuli^35, 36^. Considering their close association with ECM remodelling, we examined whether dominant murine CLPs Ym1 and Ym2 were expressed in IL-13 or IL-17a-deficient mice despite dysregulated allergic inflammation. Ym1^+^ cells in the parenchyma were evident in C57BL/6 PBS mice, likely corresponding to alveolar macrophages^41^, but as expected from previous findings where expression of Ym2 is limited in the steady state^37^, no Ym2^+^ cells were visualised in these control mice (**Fig 3a**). Chronic DRA administration in C57BL/6 wild-type mice increased the numbers of Ym1^+^ cells in the parenchyma, with Ym2 expression increased largely in the epithelium (**Fig 3a-3d**) but also in cells surrounding the airways (**Sup Fig 1c**). Whilst some cells co-expressed Ym1 and Ym2, Ym2 expression dominated in airway epithelial cells and Ym1 in parenchymal cells (**Sup Fig 1c**). Quantification of staining intensity of Ym1 and Ym2 protein revealed no significant changes between wild-type and IL-17a-defienct mice, although airway epithelial Ym2 staining was reduced (**Fig 3b**). Distribution of Ym1 and Ym2 positive cells in the lungs revealed a similar expression pattern between wild-type and IL-13-deficient mice (**Fig 3c**), although there was a small but significant reduction in levels of Ym2 in the airway epithelium in *Il13*^−/−^ compared to *Il13*^+/+^ mice (**Fig 3d**). Similarly, at the transcript level, no significant differences in mRNA expression of *Chil1*, *Chil3* or *Chil4* in whole lung were detected between wild-type and IL-17a-deficient mice (**Fig 3e**). A small but significant reduction in expression of *Chil1* and *Chil4,* but not *Chil3*, was observed in the lungs of IL-13-deficient mice compared to wild-type animals (**Fig 3d**). However, *Chil4* mRNA was still markedly increased in *Il13*^−/−^ DRA versus PBS mice.

**Fig 3:**
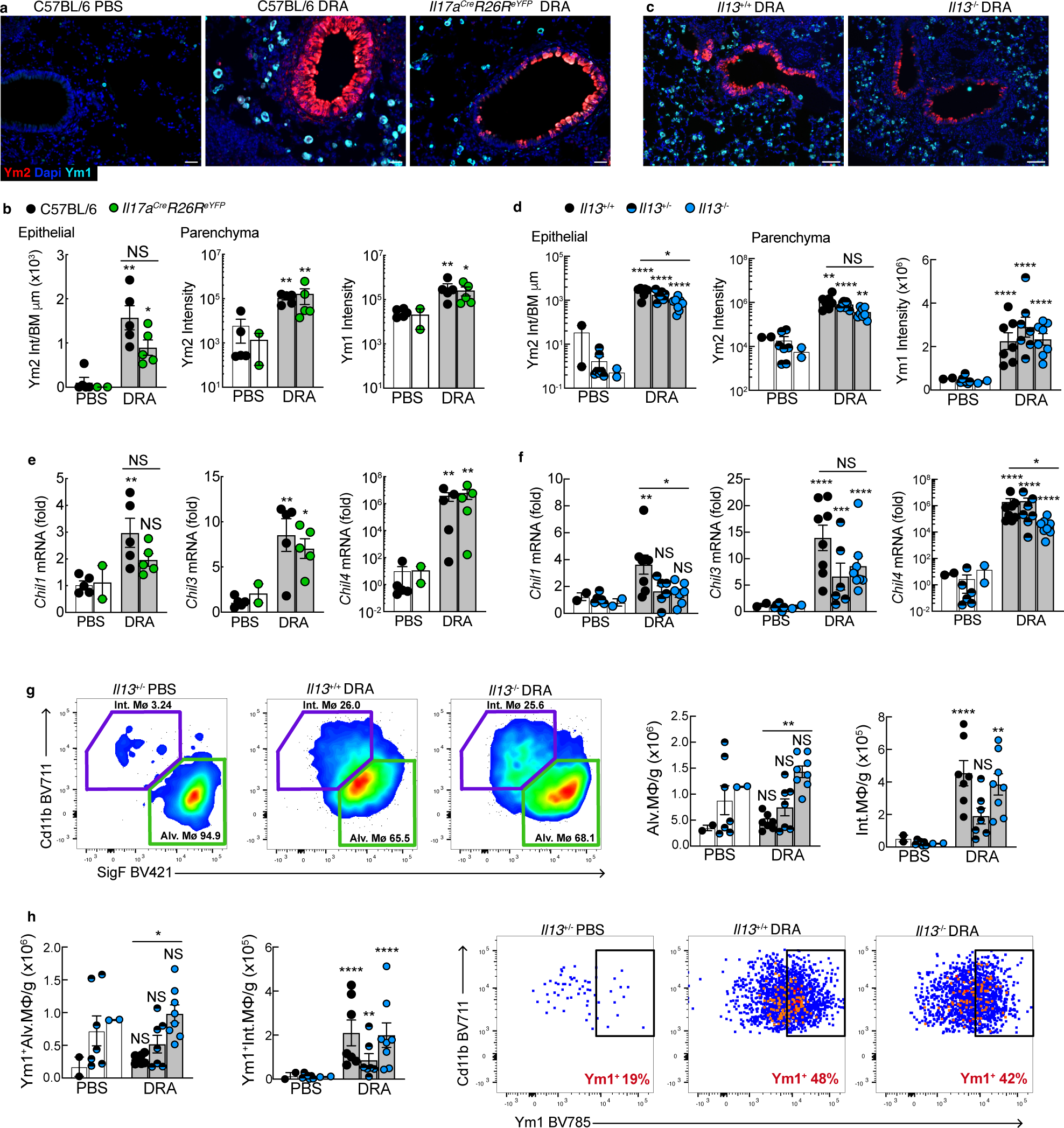
Ym1 and Ym2 are increased during allergic pathology even when inflammation is reduced. **a)** Representative images of lung tissue sections showing an airway from C57BL/6 or *Il17a*^Cre^*Rosa26*^eYFP^ female mice treated with PBS or DRA twice weekly for 8 weeks. Lung sections were stained with DAPI to visualise cell nuclei and antibodies recognising Ym1 and Ym2; scale bar=30μm. **b)** Analysis of staining intensity of Ym2 within airway epithelial cells normalised to airway basement membrane length or Ym2 and Ym2 staining intensity within regions of the lung parenchyma that exclude airways and large blood vessels from mice in **a)**. **c)** Representative images of lung tissue sections showing an airway from female *Il13*^+/+^ or *Il13*^−/−^ mice treated with DRA twice weekly for 8 weeks. Lung sections were stained as in a) with scale bar=30μm. **d)** Ym1 and Ym2 staining intensity from from mice in **c)** analysed as in **b)**. mRNA expression of *Chil1, Chil3 and Chil4* in whole lungs from **e)** C57BL/6 or *Il17a*^Cre^*Rosa26*^eYFP^ mice treated as in **a)** or **f)** *Il13*^eGFP^ wild-type, heterozygote or homozygote female mice treated with PBS or DRA twice weekly for 8 weeks. mRNA was relative to geometric mean of housekeeping genes *Gapdh*, *Rpl13a* and *Rn45s*. **g)** Representative flow cytometric identification of alveolar (Alv.Mo) and interstitial macrophages (Int.MØ) in *Il13*^eGFP^ mice treated as in **f)**. Gating based on Lineage^−^ Ly6G^−^ CD64^+^ MertK^+^ cells within whole lung tissue digests with graphs showing the number of cells per gram of lung. **h)** The total numbers of numbers of Ym1^+^ alveolar (Alv.MØ) and interstitial macrophages (Int.MØ) per g of lung in *Il13*^eGFP^ mice treated as in **f)** and analysed by flow cytometry with representative flow cytometric plots show the percentage of Ym1^+^ or Ym1^−^ interstitial macrophages (Int.MØ). Datapoints show individual animals with bars representing mean ± s.e.m with n=2-9 female mice per group and images and flow plots are representative of n=7-8 mice per group. Data are from (**a,b** and **e**) one individual experiments or (**c, d, f-h**) two combined experiments and were analysed by ANOVA with Tukey’s multiple comparison test and significance level shown relative to PBS C57BL/6 or *Il13*^+/−^ mice or between C57BL/6 and *Il17a*^Cre^*Rosa26*^eYFP^ DRA mice or *Il13*^+/+^ and *Il13*^−/−^ DRA mice as indicated on the graph. \**P*<0.05, ** *P*<0.01, *** *P*<0.001, **** *P*<0.0001 and NS, not significant.

As macrophages constitute a major source of Ym1 in the lungs during inflammation and pathology, we also examined numbers of alveolar (Alv.MØ) and interstitial macrophages (Int.MØ). Alv.MØ dominant in the lungs of PBS treated mice, with Int.MØ making up only a small proportion of the total lung Cd64^+^ MerTK^+^ macrophage pool (**Fig 3g**). After DRA administration, the proportion and numbers of Int.MØ increase in all mice. Similarly, Ym1^+^ Alv.MØ only increased in *Il13*^−/−^ DRA mice, whereas both the number and proportion of Ym1^+^ Int.MØ increase in all allergic animals regardless of IL-13 expression (**Fig 3h**). Collectively, these results demonstrate that CLPs Ym1 and Ym2 are highly upregulated in the lungs during chronic allergen administration, even in the absence of IL-13 or IL-17a signalling.

### Ym2 secretion but not Brp39 is required for allergic airway remodelling

Brp39 is the mouse homologue of YKL40 in humans. As such, it has been widely studied in models of inflammation and pathology and suggested to be required for optimal allergen sensitisation and type 2 inflammation^57^. Whilst Brp39 is increased following DRA administration^37^, *Chil1* expression was significantly reduced in allergic IL-13-deficient mice (**Fig 3f**) that exhibit airway remodelling suggesting Brp39 may not be required for excess ECM accumulation. To test this, we administered DRA to Brp39-deficient BALB/c or wild-type mice and assessed inflammation and extent of collagen deposition. Of note, BALB/c mice develop a more rapid inflammatory response to DRA compared to C57BL/6 mice^37^. However, at chronic time points, numbers of eosinophils and neutrophils are equivalent between strains, as is the magnitude of airway remodelling. Airway eosinophil and neutrophil numbers (**Sup Fig 2a**) and Th2 and Th17 cells in the lung (**Sup Fig 2b**) were increased to similar degree in both Brp39-deficient and wild-type DRA mice. Additionally, the extent of collagen deposition around the airway of allergic mice was also not regulated by Brp39, further suggesting this CLP is not key for driving ECM remodelling.

Ym1 and Ym2 are highly homologous proteins, often grouped together as one entity, Ym1/2. To this extent, very little is known about the function of Ym2 in health or disease. Ym2 is the most upregulated CLP in the lungs following allergic airway inflammation and pathology^37^ and was still significantly increased in both IL-17a and IL-13-deficient animals (**Fig 3a-f**). Therefore, we sought to determine for the first time, the role of Ym2 during allergic airway disease. We utilised CRISPR-Cas9 gene deletion approach to knock out the *Chil4* gene. Whilst sequencing the founder mice we identified a duplication of the region where the CLPs are located (**Sup Fig 3a and 3b**), as has been described elsewhere recently^58^. We determined that only one copy of *Chil4* was deleted in our founders leading to a partial knock-down of *Chil4* (**Sup Fig 3a**), hereby referred to as *Chil4*^KD/KD^ (**Sup Fig 3c and 3d**). Analysis of lung mRNA levels in allergic mice confirmed significantly reduced but not absent expression of *Chil4* in the DRA *Chil4*^KD/KD^ mice (**Fig 4a**). Whilst *Chil4*^KD/KD^ mice had normal expression of *Chil1* and *Chia* in the lungs of mice administered either PBS or DRA, there was a reduction in *Chil3* expression in allergic but not PBS animals (**Fig 4a**). Ym2 is highly secreted into the airways during pathology, and DRA *Chil4*^KD/KD^ mice had markedly reduced Ym2 levels in the BAL compared to *Chil4*^WT/WT^ (**Fig 4b**). Decreased Ym2 secretion appeared to correlate with reduced Ym2^+^ cells in the lung parenchyma but not the airway epithelium (**Fig 4c**). Similar to mRNA levels (**Fig 4a**), expression of Ym1^+^ cells in the parenchyma of the lungs were significantly decreased in *Chil4*^KD/KD^ versus *Chil4*^WT/WT^ DRA mice (**Fig 4c**). Alterations to Ym1^+^ and Ym2^+^ cells were not reflected in an overall change in the numbers of Alv.MØ, a major producer of Ym1, in the lungs of allergic animals (**Fig 4d**). However, there was a small but significant decrease in Int.MØ in *Chil4*^KD/KD^ compared to wild-type allergic mice (**Fig 4d, Fig Sup 3e**). All Alv.MØ express Ym1 in the steady state (**Fig 4e; left**), but the number of Ym1^+^ Alv.MØ are reduced upon allergic inflammation (**Fig 4e; right**) because macrophages start to secrete Ym1^41^. Knock-down of Ym2 had no impact on Ym1^+^ Alv.MØ in allergic mice but both the number and percentage of Ym1^+^ Int.MØ were significantly reduced in DRA *Chil4*^KD/KD^ mice compared to DRA wild-type animals (**Fig 4f**). Overall, we demonstrate for the first time, the generation of a Chil4 transgenic mouse, *Chil4*^KD/KD^, that exhibits reduced secretion of Ym2 into the airways and reduced Ym1^+^ and Ym2^+^ cells in the parenchyma. These changes may in part result from fewer Int.MØ accumulating in the lungs during allergic pathology.

**Fig 4:**
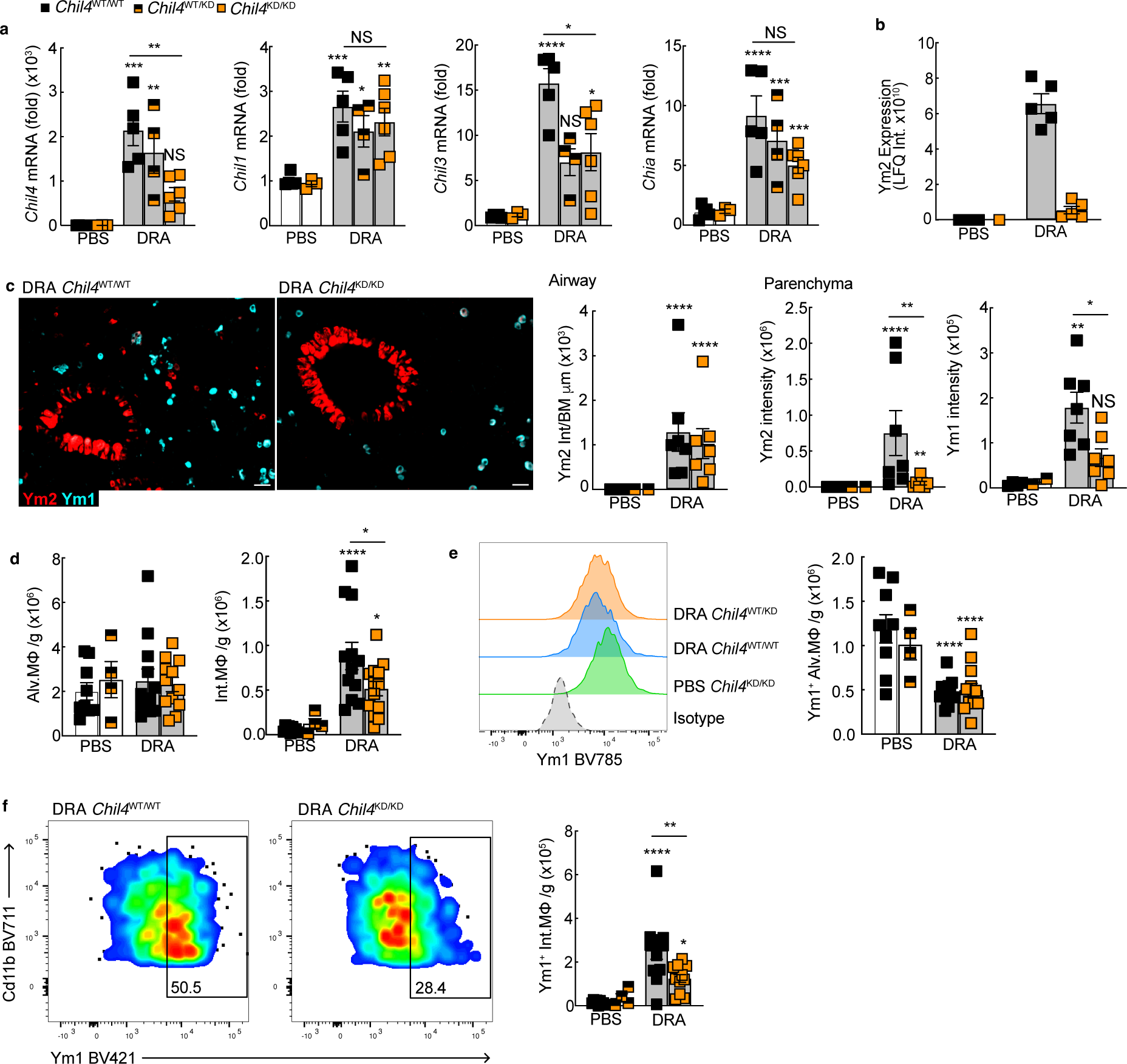
CRISPR targeting of Chil4 results in mice with reduce Ym2 secretion in the BAL. a) mRNA expression of *Chil4, Chil1, Chil3 and Chia* in whole lungs from littermate *Chil4*^KD^ mice treated with PBS or DRA twice weekly for 8 weeks. mRNA expression was relative to geometric mean of housekeeping genes *Gapdh*, *Rpl13a* and *Rn45s.* b) Secreted Ym2 levels in the BAL fluid of mice as in a) analysed by mass spectrometry; LFQ label free quantitation. c) Representative images of lung tissue sections from *Chil4*^WT/WT^ and *Chil4*^KD/KD^ mice treated with DRA as in a), stained with antibodies recognising Ym1 and Ym2 (scale bar=30μm). Graphs show analysis of staining intensity of Ym2 within airway epithelial cells normalised to airway basement membrane length or Ym1 and Ym2 staining intensity within regions of the lung parenchyma. d) Numbers of alveolar (Alv.MØ) and interstitial macrophages (Int.MØ) in whole lung tissue from mice as in a) analysed by flow cytometry. e) Representative histograms showing intracellular expression of Ym1 in Cd64^+^ MertK^+^ SigF^+^ Cd11c^+^ Alv.MØ compared to isotype stained cells from mice as in a). Graph shows numbers of Ym1^+^ Alv.MØ per g of lung tissue. f) Representative flow cytometry plots of intracellular Ym1 expression in Int.MØ from mice treated as in a). Gates based on isotype staining for intracellular Ym1 and numbers show percentage of Ym1^+^ Int.MØ. Graph shows numbers of Ym1^+^ Int.MØ per g of lung tissue. Data for a-c are representative from 2 individual experiments (n=3-6 per group) and d-f from two combined experiments (n=4-12 female mice per group). Data were analysed by ANOVA with Tukey’s multiple comparison test and significance level shown relative to PBS *Chil4*^WT/WT^ mice or between *Chil4*^WT/WT^ and *Chil4*^KD/KD^ DRA mice as indicated on the graph. \**P*<0.05, ** *P*<0.01, *** *P*<0.001, **** *P*<0.0001 and NS, not significant.

Allergen-induced neutrophil or eosinophil recruitment was not altered in *Chil4*^KD/KD^ mice (**Fig 5a**) and there were no overt differences in T cells, ɣδT cells or ILC populations (**Fig 5b and Sup Fig 3f**) or IL-17a and IFNɣ expressing T cells compared to wild-type mice (**Fig 5c**, **5d**). However, DRA mice that failed to secrete Ym2 had significantly reduced numbers of IL-4^+^ CD4^+^ T cells (**Fig 5d**) and reduced *Il5* mRNA expression in the lungs (**Fig 5e**). To determine whether the small reduction in IL-4 and IL-5 in *Chil4*^KD/KD^ mice cumulated in a skewed inflammatory response, levels of RELM⍺ (Retnla), a hallmark of a strongly polarised type 2 immune response, were measured in the lungs. Whilst *Retnla* expression did not change in *Chil4*^KD/KD^ versus *Chil4*^WT/WT^ allergic mice (**Fig 5e**), there were significantly fewer RELM⍺^+^ Alv.MØ and Int.MØ compared to allergic wild-type mice (**Fig 5f and Sup Fig 3g, 3h**). To further broadly investigate differences in immune responses between wild-type and *Chil4*^KD/KD^ mice, we performed differential gene expression analysis of whole lung RNA after 8 weeks of DRA administration using the Nanostring nCounter Myeloid Innate Immunity Panel. Coinciding with a decrease in IL-4^+^ CD4^+^ T cells, genes associated with type 2 immunity such as *Ccl24, Retnla, Arg1, Mmp12* were significantly reduced in allergic *Chil4*^KD/KD^ compared to *Chil4*^WT/WT^ mice (**Sup Fig 4a**). Analysis also revealed over-representation of genes within the Th2 activation and differentiation and maintenance of myeloid cells pathways in *Chil4*^KD/KD^ allergic mice (**Sup Fig 4b**).

**Fig 5:**
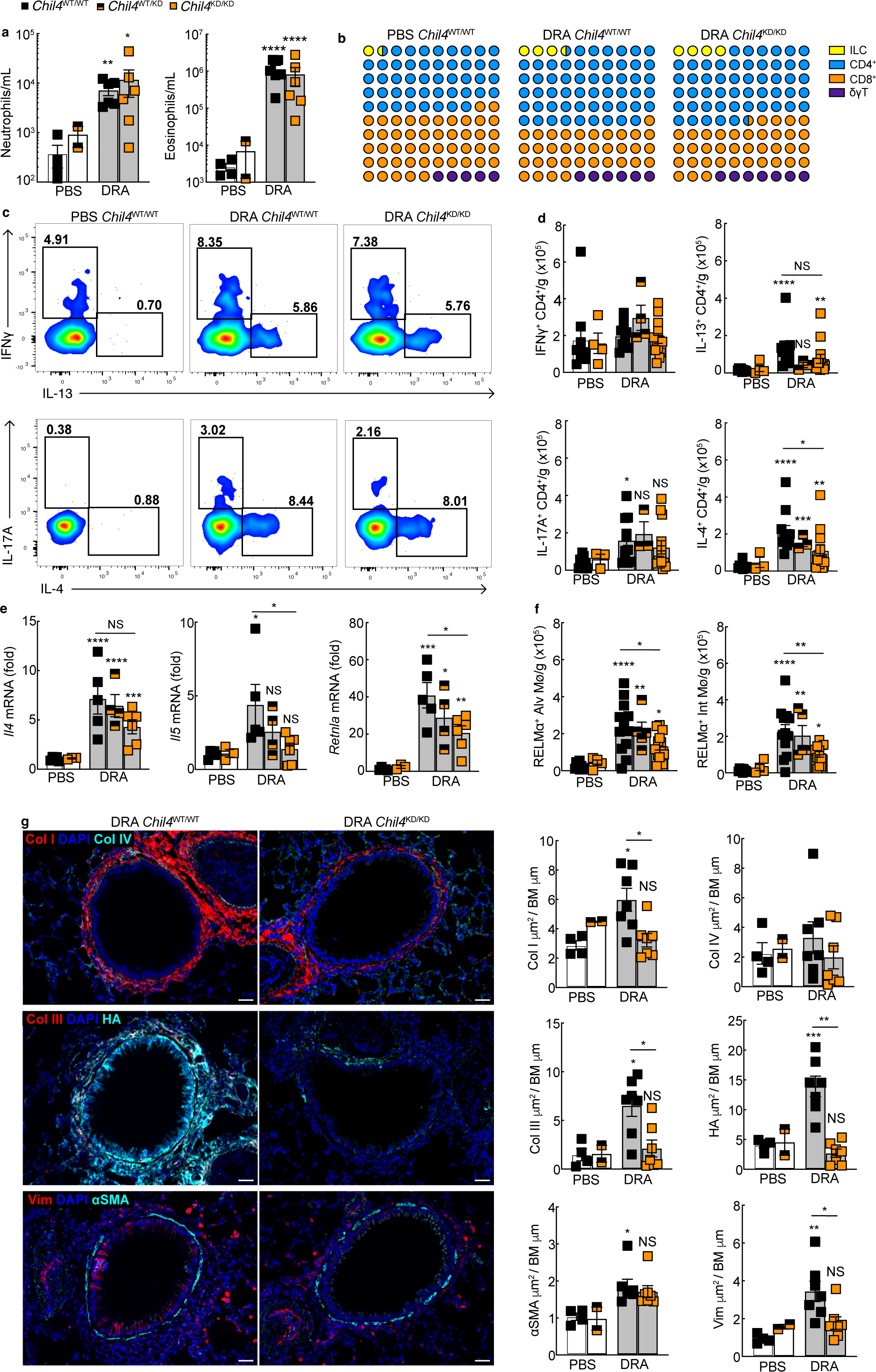
Knock down of Ym2 prevents allergen-induced accumulation of ECM. **a)** *Chil4*^KD^ littermate mice were treated with PBS or DRA twice weekly for 8 weeks. Numbers of neutrophils and eosinophils in the BAL of mice were analysed by flow cytometry. **b)** Waffle plot showing relative proportions of Lineage-CD90^+^ CD25^+^ innate lymphoid cells, CD4^+^ TCRβ^+^, CD8^+^ TCRβ^+^, TCRƔδ^+^ TCRβ^−^ cells in the lungs of mice in **a)** analysed by flow cytometry. **c)** Representative flow cytometry plots showing intracellular cytokine expression in CD4^+^ TCRβ^+^ cells from single cell suspensions of lungs from mice in a) stimulated ex vivo with PMA/ionomycin prior to staining with antibodies. Gates are based on cells stained with an isotype and numbers represents the cytokine positive cells as a percentage of all CD4^+^ TCRβ^+^ cells. **d)** Analysis of flow cytometry data in **c)** was used to calculate numbers of cytokine positive CD4^+^ TCRβ^+^ cells per g of lung tissue. **e)** mRNA expression of *Il4, Il5,* and *Retnla* in whole lungs from littermate *Chil4*^KD^ mice treated as in **a)**. mRNA was relative to geometric mean of housekeeping genes *Gapdh*, *Rpl13a* and *Rn45s*. **f)** Numbers of alveolar (Alv.MØ) and interstitial macrophages (Int.MØ) expressing intracellular RELM⍺ in single cell lung suspensions from mice as in a) analysed by flow cytometry. **g)** Representative images of lung tissue sections from *Chil4*^WT/WT^ or *Chil4*^KD/KD^ mice treated with DRA for 8 weeks and then stained with DAPI to visualise cell nuclei and antibodies or binding proteins recognising Collagen I and IV; collagen III and HA binding protein (HABP); or ⍺SMA and vimentin (scale bar = 30μm). Graphs show analysis of positive-stained area around the airways for each antigen and normalised to length of basement membrane for *Chil4*^KD^ littermate mice treated with PBS or DRA for 8 weeks. Data for **b-d** and **f**, are from two combined experiments (n=4-12 female mice per group) and data from **a, e,** and **g** are representative from 2 individual experiments (n=2-7 female mice per group). Datapoints show individual animals with bars representing mean ± s.e.m. Data were analysed by ANOVA with Tukey’s multiple comparison test and significance level shown relative to PBS *Chil4*^WT/WT^ mice or between *Chil4*^WT/WT^ and *Chil4*^KD/KD^ DRA mice as indicated on the graph. \**P*<0.05, ** *P*<0.01, *** *P*<0.001, **** *P*<0.0001 and NS, not significant.

DRA allergen administration led to the expected increase in total collagen, goblet cell hyperplasia, airway ECM, airway smooth muscle mass and vimentin^+^ cells (**Sup Fig 3i and Fig 5g**). However, in mice where Ym2 secretion was impaired, DRA-induced ECM accumulation was significantly reduced compared to wild-type mice. Remarkably, expression of collagen I, III and HA around the airways in *Chil4*^KD/KD^ mice remained at levels observed in non-allergic animals (**Fig 5g**). There was also a significant reduction in the number of vimentin^+^ cells surrounding the airway in *Chil4*^KD/KD^ mice compared to *Chil4*^WT/WT^. However, airway muscle mass (**Fig 5g**) and PAS^+^ cells (**Fig Sup 3i**) were increased in all mice exposed to DRA, despite a small reduction in type 2 immune responses (**Fig 5c-e and Sup Fig 4**). All together our data demonstrates a key pathogenic role for Ym2 during chronic allergic pathology, whereby secreted Ym2 contributes to type 2 immune response and is required for remodelling of the airway matrix.

### Ym2 secretion enhances collagen-crosslinking

We next aimed to gain a deeper understanding of the collagen networks that develop during chronic allergic pathology and how Ym2 might be regulating this matrix. Lung tissue sections from *Chil4* littermate mice treated with PBS or DRA for 8 weeks were stained with picrosirius red and imaged with polarised light to examine birefringence of collagen fibres (**Fig 6a**). Under polarised light, thin less organised collagen fibres display greenish colours whilst thicker organised bundles of mature fibre networks display yellow and red/orange colours^59, 60^. Following DRA exposure, there was a reduction in the overall amount of collagen around the airways of *Chil4*^KD/KD^ versus *Chil4*^WT/WT^ animals (**Fig 6a**, **6b**), and more specifically fibres that were red/orange and yellow (**Fig 6b**). Additionally, the proportion of thicker bundles of collagen (red/orange and yellow fibres) were reduced in *Chil4*^KD/KD^ compared to *Chil4*^WT/WT^ allergic mice with an increase in thinner, less organised fibres (green) when DRA was administered for 8 weeks (**Fig 6c**). Formation of mature organised collagen fibres relies on intermolecular cross-linking of collagen fibrils by enzymes such as lysyl oxidases (Lox). Dysregulation of Lox family members have been highlighted in numerous fibrotic conditions^61^ and more recently in ECM remodelling in asthma^62^. To ascertain whether CLPs might be driving ECM remodelling, in part through regulation of collagen cross-linking, we administered DRA to *Chil4* littermate mice for 2 weeks (**Fig 6d**). Two weeks of allergen exposure was already enough to observe signs of collagen deposition around the airways of *Chil4*^WT/WT^ but not *Chil4*^KD/KD^ mice (**Fig 6e-g**). During this early stage of ECM remodelling, there was an increase in expression of Lox specifically in the cells of the lung parenchyma surrounding the airways of *Chil4*^WT/WT^ DRA versus PBS mice (**Fig 6f and 6h**). However, DRA administration failed to upregulate Lox protein levels in the lung parenchyma of *Chil4*^KD/KD^ mice. Additionally, the expression of Lox in the airway epithelium also remained significantly lower in *Chil4*^KD/KD^ versus *Chil4*^WT/WT^ DRA animals. As IL-13-deficient animals still exhibit normal collagen remodelling (**Fig 3**) we hypothesised IL-13 would not regulate *Lox* expression. Administration of DRA to *Il13*^−/−^ mice for 2 weeks revealed no change in total airway collagen content compared to DRA mice (**Fig Sup 5a**). Goblet cell hyperplasia did not develop in *Il13*^−/−^ DRA mice as expected (**Fig Sup 5b**). Furthermore, there was no significant change in Ym1 or Ym2 levels in the lungs of allergic IL-13-deficient compared to wild-type mice (**Fig Sup 5c**). Expression of *Lox* in whole lung tissue was not different in *Il13*^+/+^ versus *Il13*^−/−^ mice (**Fig Sup 5d**). Together data demonstrates that Ym2 increases Lox expression in the lung during allergic airway pathology, which may be one mechanism through which Ym2 contributes to lung ECM remodelling.

**Fig 6:**
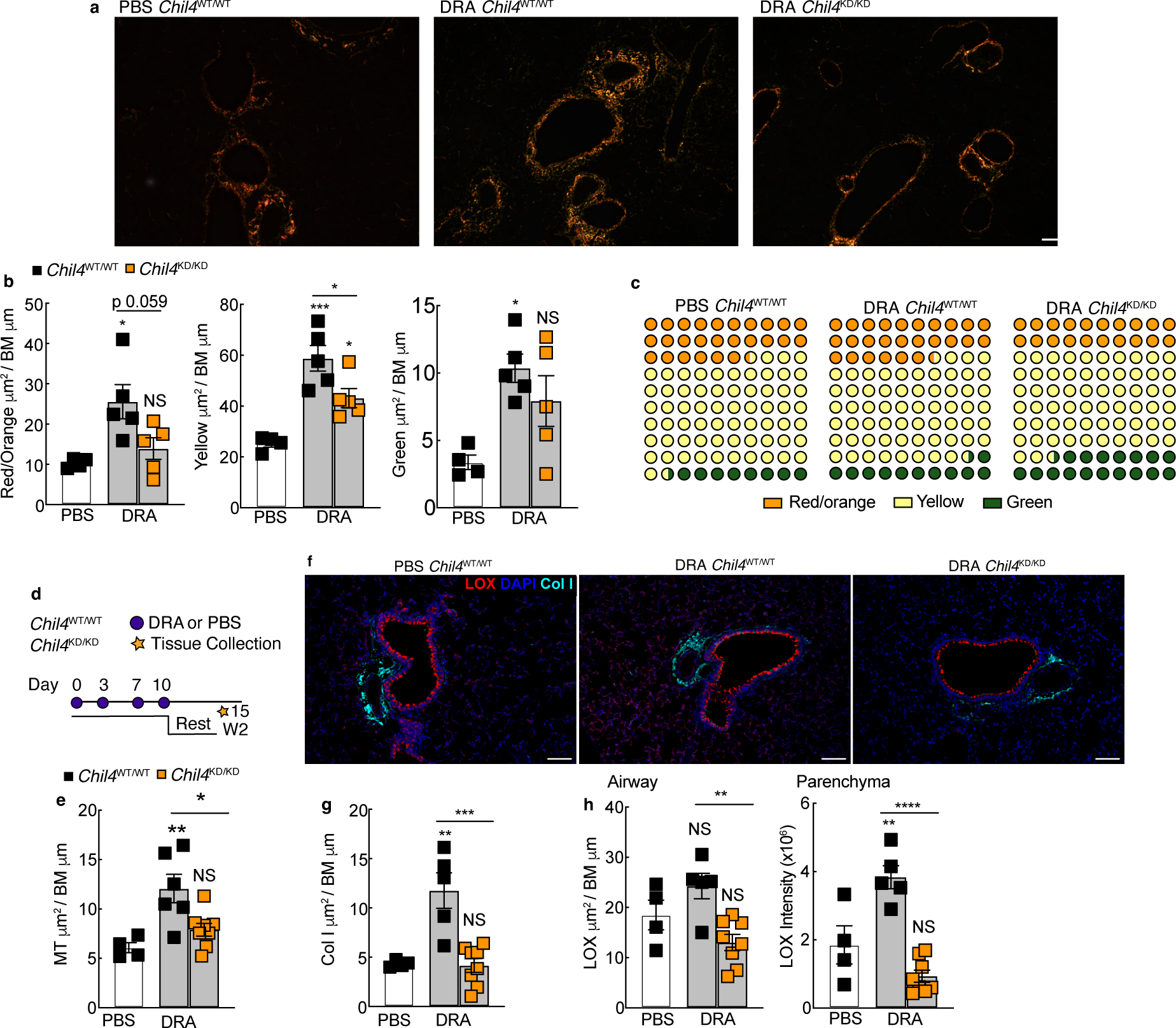
Ym2 regulates lysyl oxidase and collagen organisation in the lungs during allergic pathology. Lung sections from *Chil4*^KD^ littermate female mice treated with PBS or DRA for 8 weeks were stained with picrosirius red and images captured under polarised light. **a)** Representative images of picrosirius red staining under polarised light (scale bar =200μm) with **b)** graphs showing analysis of area around the airways of lungs for birefringence collagen separated into red/orange, yellow and green fibres normalised to basement membrane length for each airway. **c)** Waffle plots show proportion of birefringence collagen fibres that were red/orange, yellow or green. **d)** *Chil4*^KD^ littermate mice were treated with PBS or DRA twice weekly for 2 weeks and lung tissue collected 5 days after the last PBS or DRA dose. **e)** Histological analysis for Masson’s trichrome (MT) stained tissue sections PBS and DRA *Chil4*^KD^ mice treated as per **d)**. **f)** Representative images of lung tissue sections from PBS or DRA *Chil4*^WT/WT^ and DRA *Chil4*^KD/KD^ mice treated as in **a)** stained with DAPI to visualise cell nuclei and Lox and Collagen I (scale bar = 100μm). Analysis of positive-stained area around the airways normalised to length of basement membrane for **g)** collagen I and **h**) Lox. Intensity of Lox staining within the lung parenchyma and for *Chil4*^KD^ littermate mice treated as in **a)** was also analysed. Data for a-c are representative from 2 individual experiments (n=3-6 female mice per group) and d-h from one individual experiment (n=4-8 male and female mice per group). Datapoints show individual animals with bars representing mean ± s.e.m. Data were analysed by ANOVA with Tukey’s multiple comparison test and significance level shown relative to PBS *Chil4*^WT/WT^ mice or between *Chil4*^WT/WT^ and *Chil4*^KD/KD^ DRA mice as indicated on the graph. \**P*<0.05, ** *P*<0.01, *** *P*<0.001, **** *P*<0.0001 and NS, not significant.

### Therapeutic inhibition of Ym1 reverses ECM remodelling

Ym2 was key for driving pathogenic ECM remodelling (**Fig 5 and 6**). However, *Chil4*^KD/KD^ allergic mice also have reduced *Chil3* mRNA expression together with Ym1^+^ cells in the lungs (**Fig 4a and c**). Combined with previous reports suggesting a role for Ym1 in tissue repair and fibrosis^41, 63^, we wanted to determine whether ECM remodelling required sustained expression of CLPs. ECM remodelling was allowed to develop in wild-type BALB/c mice by administration of DRA for 4 weeks^37^. Mice were then given an additional 4 weeks of DRA (or PBS) alongside administration of an antibody that would neutralise Ym1^41, 46, 64^ (**Fig 7a**). Similar to *Chil4*^KD/KD^ mice, blocking Ym1 had no impact on chronic neutrophil or eosinophil inflammation (**Fig 7b**), nor was there significant changes to T cells, cytokine production (**Fig 7c**) or PAS^+^ epithelial cells (**Fig 7d**). Despite this, anti-Ym1 treatment reversed the allergen-induced collagen accumulated around the airways (**Fig 7e**). To determine whether the effect on collagen might be explained by changes in Ym2, expression of Ym2^+^ cells were examined in the lungs. Neutralisation of Ym1 in allergic mice significantly reduced not only Ym1 immuno-staining but also Ym2 expression in the lung parenchyma (**Fig 7f**). However, no significant change in Ym2 was observed in the airway epithelium of anti-Ym1 versus IgG2a treated allergic animals (**Fig 7f**). We next investigated whether blocking Ym1 was regulating specific collagen subtypes by examining expression of airway collagen I, IV and III. As previously shown following DRA exposure in BALB/c mice, collagen I levels, rather than collagen III, appears to dominate the total increase in collagen around the airway^37^ (**Fig 7g**). However, anti-Ym1 treatment significantly reduced collagen I accumulation to levels found in PBS challenged mice. Airway collagen IV expression decreased after allergen exposure, and this was not rescued by anti-Ym1 treatment (**Fig 7g**). The amount of HA around the airways was also completely reduced to levels found in non-allergic animals by anti-Ym1 treatment, as were allergen-induced increases in airway smooth muscle and vimentin expressing cells (**Fig 7g**). Overall, these results suggest that Ym1 is required to not only sustain ECM remodelling but that blocking Ym1 can initiate a process that leads to degradation of any ECM molecules already deposited around the airways. Thus, neutralisation of a CLP has the capacity to reverse allergic airway remodelling.

**Fig 7:**
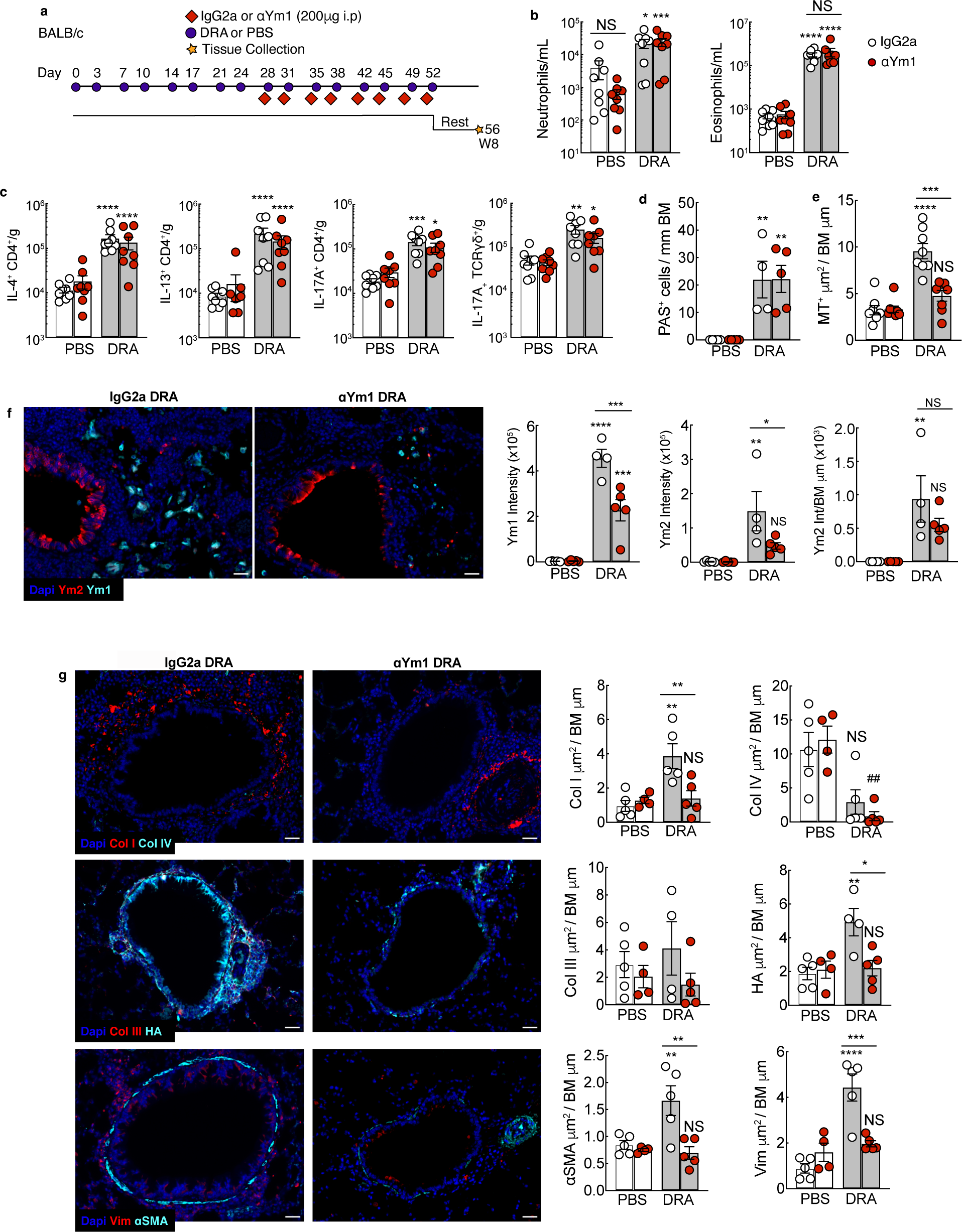
Therapeutic neutralisation of Ym1 reverses allergen-induced ECM accumulation. **a)** BALB/c mice were treated with PBS and DRA intranasally twice weekly for 8 weeks. At the 4 week time point mice were also treated intraperitoneally with either IgG2a or anti-Ym1 for 4 weeks. **b)** Numbers of neutrophils and eosinophils in the BAL of mice were analysed by flow cytometry. **c**) Numbers of CD4^+^ TCRβ^+^, cells expressing intracellular cytokine IL-4, IL-13 or IL-17a and TCRƔδ^+^ TCRβ^−^ cells expressing intracellular IL-17A from single cell suspensions of lungs from mice in **a)** stimulated ex vivo with PMA/ionomycin prior to staining with antibodies and performing flow cytometry. **d)** Lung tissue sections were stained with PAS and numbers of PAS^+^ airway epithelial cells counted per airway and normalised to length of the basement membrane from mice as in **a**). **e**) Lung tissue sections were stained with Masson’s trichrome and MT^+^ area was analysed around the airway excluding vascular regions and normalised to basement membrane length of mice as in **a**). **f**) Representative images of lung tissue sections showing from mice as in **a**) stained with DAPI to visualise cell nuclei and antibodies recognising Ym1 and Ym2 (scale bar=30μm). Graph shows analysis of staining intensity of Ym1 and Ym2 in the lung parenchyma and Ym2 within airway epithelial cells normalised to airway basement membrane length. **g**) Representative images of lung tissue sections from BALB/c mice treated as in a) stained with DAPI to visualise cell nuclei and antibodies or binding proteins recognising ⍺SMA, vimentin or Collagen I and IV or collagen III and HA binding protein (HABP) (scale bar = 30μm). Graphs show analysis of positive-stained area around the airways for each antigen and normalised to length of basement membrane. Data are from b, c and e are pooled from 2 individual experiments (n=8 per group) and data from **d, f** and **g** are representative of 2 individual experiments (n=5 per group). Datapoints show individual animals with bars representing mean ± s.e.m. Data were analysed by ANOVA with Tukey’s multiple comparison test and significance level shown relative to PBS IgG BALB/c mice or between IgG treated DRA mice and anti-Ym1 DRA treated mice as indicated on the graph. \**P*<0.05, ** *P*<0.01, *** *P*<0.001, **** *P*<0.0001 and NS, not significant.

## Discussion

Our studies described here, demonstrate a clear requirement for the CLPs Ym1 and Ym2 in the development and maintenance of ECM remodelling during allergic airway pathology. The study of CLPs has been clouded by limited knowledge of CLP biology with little known about their receptors or respective signalling pathways. Understanding the role of CLPs in regulating airway remodelling is also obfuscated by the fact that these molecules can be strongly induced by classical IL-4/13 signalling pathways in many conditions^41, 65–67^, a signalling pathway synonymous with tissue repair and fibrosis. Due to their association with type 2 immunity and reported abilities to interact with GAGs^38, 40^ and collagen^39^, a role for CLPs in tissue remodelling has been assumed. Data shown here, combined with our previous studies demonstrating that Ym1 can promote lung tissue repair independently of IL-4R⍺ signalling^41^, now provide strong evidence that CLPs can drive ECM remodelling. In humans YKL40, which shares functional similarities with Ym1 in mice, has been shown to bind collagen I in biochemical assays regulating fibril formation^39^, and to prevent collagen degradation^68^ and enhance collagen output from fibroblasts^69^ using *in vitro* assays. Here, we demonstrate Ym2 enhancement of Lox expression in the lungs. Lysyl oxidase is an enzyme necessary for the assembly and strengthening of fibrillar collagen and elastic fibres^70^ through regulation of ECM cross-linking. Inhibition of the LOX family of enzymes can abrogate fibrotic pathologies^71–73^ and alter cell migration through the matrix^74^ with the potential to influence immune responses. Disruption of the CLPs upstream of LOX may be an attractive avenue to not only broadly influence the ECM microenvironment, but also fine-tune the immune response in order to promote tissue health. Historically it has been very difficult to study the ECM as knock-out animals often have severe or even lethal phenotypes (e.g. ^75, 76^). The possibility of interrogating the downstream effects of remodelling using CLP transgenic mice could help disentangle the roles of inflammation, tissue remodelling, and extracellular matrix composition not only in allergic airway pathology but also many other disorders that feature pathogenic ECM remodelling. A significant finding from this study comes from our attempts to block the function of these CLPs after the initiation of allergic remodelling. Not only is remodelling prevented in the absence of secreted Ym2, but neutralising Ym1was able to reverse remodelling after it had become established. The requirement for sustained CLP expression in progressive remodelling has interesting implications not only for potential treatments but also for understanding the process of remodelling itself and the specific role of these proteins in a range of chronic conditions. Increased levels of Ym1 in mice, as well as YKL40 in humans have been observed in many different pathologies^35, 36, 56^, often in local tissue environments as well as in the serum.

The entire CLP genomic locus is a hotspot for genetic duplications, as evidenced by the recent duplication of *Chia1*, *Chil3*, and *Chil4* in the C57BL/6J mouse^58, 77^. These different gene products are likely to have redundant functions that cloud their individual roles, much in the same way that IL-4 can partially compensate for IL-13 in certain circumstances^78, 79^. One possibility is that a balance between different CLPs and the total combined levels of these mediators determines disease outcome. This would fit with our data showing that remodelling is reduced in *Chil4*^KD^ mice which have reduced levels of multiple CLPs after induction of allergic pathology. *Chil4*^KD^ mice still express Ym2 but appear not to secrete it into the airway lumen, which also hints at the importance of CLP localisation for effector function. Despite high sequence homology between CLPs, their distinct expression patterns in the naïve and allergic lung^37^, suggest CLPs are eliciting different responses in these settings. Thus, the balance between levels of Ym1 and Ym2, or CLPs in general, could be a key determinant of the extent of tissue pathology and ECM remodelling.

Despite recognition of its clinical importance, airway remodelling is still a relatively understudied area of asthma research. Whilst inflammation can contribute towards features of remodelling, clinical data now recognises that airway remodelling is not purely a downstream consequence of immune cells and mediators^29, 80–82^. IL-13 and IL-17a are traditionally thought of as major mediators of allergic inflammation^16, 83–85^, with IL-13 also a key regulator of epithelial remodelling processes such as mucus secretion^54, 86^. However, using a mouse model that recapitulates some of the features of severe asthma pathology, we show that ablation of IL-13 or IL-17a had little effect on airway remodelling, with Il-13 regulating only goblet cell hyperplasia and accumulation of HA. These results were somewhat surprising considering IL-13, in various models, has been shown to also modulate specific components of the ECM^51, 87, 88^, in addition to HA^55^. Additionally, many matrix-associated proteins, such as certain matrix metalloproteinases (MMP12, MMP14) are highly driven by Th2 cytokine production^13, 89^, with classical ‘Th2’ cell types such as eosinophils also being important for producing proteins such as PRG2, which can then drive matrix changes^90^. Mice deficient for IL-13 or IL-17a and by association disruption of eosinophil/neutrophil effector immune responses were not enough to prevent some features of airway remodelling. The disconnect between IL-13/IL-17a and changes to the ECM suggests that perhaps we should treat ECM remodelling as a distinct entity from the classical allergic immune responses. These processes are likely interacting together to drive the different features of allergic airway inflammation but are clearly disparate in many aspects.

Collectively, we show that the change in magnitude and consistency of the ECM remodelling response are independent of IL-17a and IL-13 signalling, but remarkably requires functional CLP expression. To understand the roles and functions of the different aspects of airway remodelling, future studies should aim to examine how the matrix impacts the development of the other facets of disease including changes to the airway epithelium, alterations in the underlying smooth muscle and inflammatory cell recruitment^1^. For instance, ECM alterations may alter the downstream immune responses which can then in turn regulate matrix components in a feed-forward loop. Interruption of early CLP-meditated changes to the matrix could prove a potent method for regulating the immune response and long-term tissue modification^91^ in asthma and beyond. Such a treatment approach could be beneficial in cases such as paediatric asthma where signs of airway remodelling have been observed prior to disease manifestation^82, 92^, but will have limited benefits if ECM remodelling manifests as a late consequence of disease processes. It is also key to consider that blocking CLPs once a pathogenic matrix has formed, could reverse ECM remodelling. Thereby, CLP inhibitors also provide a critical avenue to explore cellular and molecular mechanisms that could be exploited for the treatment of asthma and other lung diseases.

## Methods

### Experimental animals and ethics

All animal experiments were performed in accordance with the UK Animals (Scientific Procedures) Act of 1986 under a Project License (70/8548; PP4115856) granted by the UK Home Office and approved by the University of Manchester Animal Welfare and Ethical Review Body. Euthanasia was performed by asphyxiation in a rising concentration of carbon dioxide.

Wild-type (BALB/c or C57BL/6J) mice were obtained from a commercial supplier (Envigo, Hillcrest, UK or Charles River, Margate UK). *Il13^eGFP^*(C57BL6/J)^52^, *Il17a*^Cre^*Rosa26*^eYFP^ C57BL/6 (originally provided by Dr. Brigitta Stockinger^45, 93^), *Chil1*^−/−^ BALB/c^94^ (kind gift from Dr. Alison Humbles) and *Chil4*^KD/KD^ C57BL6 mice were bred at the University of Manchester. Wild-type mice bred in house were included for all experiments involving transgenic mice. Mice used where either female, or mixed sex as indicated in the figure legend for each experiment and were between 7-14 weeks old at the start of the experiment. Animals were housed in individually ventilated cages maintained in groups of 3-6 animals in specific pathogen-free facilities at the University of Manchester. For most experiments mice were not randomised in cages, but each cage was randomly assigned to a treatment group. For experiments using *Il13*^eGFP^ and *Chil4*^KD/KD^ animals, mice were bred as littermates and were randomised in cages with investigators blind to mouse identity during necropsy. Pilot experiments were carried out with 3 mice per group to calculate sample size based on the number of animals needed for detection of a ∼35% change in Masson’s trichrome positive area around the airway at *P* value of <0.05.

### Generation of Chil4 ^KD/KD^ mice

CRISPR can generate knockout alleles by generating frameshifting Insertions or Deletions within coding regions of genes. However, due to the high sequence homology between *Chil3* and *Chil4* coding regions we were unable to identify *Chil4* specific sgRNA. Instead, we designed CRISPR sgRNA targeted to the introns flanking critical exon 5 of the *Chil4* gene intended to specifically excise this exon. The sgRNA sequences, tatgatgcttggaaaataaa, gtaacacagcacctaggtga and actattgttcttaaacagca were purchased as crRNA oligos, and were annealed with tracrRNA (Integrated DNA Technologies) in sterile, RNase free injection buffer (TrisHCl 1mM, pH 7.5, EDTA 0.1mM) by combining 2.5μg crRNA with 5μg tracrRNA and heating to 95°C, which was allowed to slowly cool to room temperature. For embryo microinjection the annealed sgRNA were each complexed with Cas9 protein (New England Biolabs) at room temperature for 10min in injection buffer (final concentrations; each sgRNA 20ng/μl, Cas9 protein 60ng/μl) before microinjection into the nuclei of C57BL/6J (Envigo) zygotes using standard protocols. Zygotes were cultured overnight, and the resulting 2 cell embryos surgically implanted into the oviduct of day 0.5 post-coitum pseudopregnant mice. Resulting mice were screened using a PCR strategy designed to amplify across *Chil4* exon 5 and identify alleles with a size change indicative of exon deletion. Primers JA02 Geno F1 (acagacggttgttatattttgctct) and JA02 Geno R1 (cccttcttcaagcccagtca) amplify a 1041bp product on unmodified alleles, and a founder with a significant deletion was observed. Sequencing of the amplicon and alignment to genomic sequences revealed this founder to have a 331bp deletion removing exon 5 and upstream splice acceptor sequences.

### Droplet Digital PCR

Due to the incomplete loss of Ym2, and a complete mouse genome hybridisation study identifying regions of gene duplication C57BL/6J vs BALB/c mice^77^, including the locus containing *Chia*, *Chil3*, and *Chil4* we used Droplet digital PCR (ddPCR) to confirm duplication in our mouse strains. ddPCR was conducted using the QX200 Droplet Digital PCR System (BioRad). Primers and probes were generated to respective CLPs (**Supplementary Table 1**). Samples were generated as recommended by the manufacturer using the ddPCR Supermix for Probes (No dUTP) (186-3023), 900mM forward and reverse primers alongside 250mM hydrolysis probes. Oil emulsion was generated using the QX200 Droplet Generator and Droplet Generation Oil for Probes (1863005), DG8 Gaskets (1863009) and DG8 Cartridges (1864008). Oil emulsions were then loaded onto ddPCR™ 96-Well Plates (12001925) and sealed with a PCR Plate Foil Heat Seal (1814045) using a PX1 PCR Plate Sealer (1814000). PCR products were then amplified using a C1000 Touch Thermalcycler (BioRad) using manufacturer suggested settings and an annealing temperature of 60°C. Amplified droplets were then measured using the QX200 Droplet Reader and results analysed using QuantaSoftTM Analysis Pro to determine the copy number of the specific genes of interest in relation to a control gene *Tfrc* (Transferrin Receptor). Of note primers for *Chil4* in the ddPCR assay bind outside exon 5, which was targeted in the CRISPR deletion. Therefore, both copies of the *Chil4* gene are detected, despite *Chil4*^KD/KD^ mice having a functional knock out for one gene copy.

### Chil4 Knockdown colony genotyping

Due to the gene duplication, the founder genotyping strategy was unable to distinguish between wild-type, heterozygous or homozygous for the *Chil4* exon deletion. Further sequence analysis revealed that one of the 5’ intronic sgRNA also edited the duplicated target sequence resulting in small 6bp deletion, which co-segregated with the exon deletion. This deletion removed an AleI restriction site in this region, allowing us to build a colony genotyping strategy based on the presence/absence of this restriction enzyme site at the deleted exon. The primers JA02 GenoF1 (ACAGACGGTTGTTATATTTTGCTCT) and JA02 LOA GenoR (ACTGACTTAATGAATGTCTGACGG) amplicons were digested with (AleI-v2, New England Biolabs). *Chil4*^KD/KD^ animals have an undigested 405bp band and wild-type animals showing two bands cleaved around 150bp and 250bp, whereas heterozygotes have all three bands (**Sup Fig 3c**).

### DRA Model of allergic airway pathology

Allergen DRA cocktail containing 5µg House **D** ust Mite (*Dermatophagoides pteronyssinus*) 5450 EU, 69.23mg per vial, 50µg **R**agweed (*Ambrosia artemisifolia*), 5µg **A** *spergillus fumigatus* extracts (Greer Laboratories) was prepared freshly for each use in vivo. As described previously, allergic airway pathology was induced by intranasal administration of 20µL of DRA cocktail or PBS to mice that were briefly anaesthetised via inhalation of isoflurane^37, 44^. DRA or PBS was administered twice weekly for up to 8 weeks and mice were rested for 5 days prior to performing BAL and collecting lung tissues.

### Isolation of cells from the BAL and lung tissue of mice

Following exsanguination, the trachea was cannulated and the airways washed four times with 0.4mL PBS. BAL was centrifuged and cells processed for flow cytometry whilst lavage fluid were stored at −20°C. Lungs were processed as previously described^37^. Briefly, a right lobe was removed and minced in 1mL of HBSS buffer containing 0.4U/mL Liberase TL (Sigma) and 80U/mL DNase type I (Thermo) for 25min in a 37°C shaking incubator. Tissue digestion was stopped by addition of 2% FBS (Gibco) and 2mM EDTA prior to passing the suspension through a 70µm cell easystrainer (Greiner Bio-one). Red blood cells were lysed (Sigma) and total live BAL and lung cells assessed with Viastain acridine orange and propidium iodide and counted using the Cellometer Auto2000 automated cell counter (Nexcelom Bioscience).

### Flow Cytometry

Equal cell numbers from lung and BAL samples for each animal were stained for flow cytometry. Cells were washed with ice-cold PBS and stained with Live/Dead Aqua or Blue (Thermo) for 10min at room temperature. All samples were then incubated with Fc block (5µg/mL CD16/CD32 (Biolegend) and 0.1% mouse serum (Sigma) in FACs buffer (PBS containing 0.5% BSA and 2mM EDTA)) for 20min prior to staining specific surface markers with fluorescence-conjugated antibodies (25min at 4°C) (Supplementary Table 2) Following surface staining, cells were fixed with ICC fix (Biolegend) and stored at 4°C until intracellular staining was performed or cells were acquired.

To measure intracellular RELM⍺ and Ym1, fixed cells were permeabilised with the Transcription Factor staining kit (eBioscience) for 20min at room temperature and stained for 45min with rabbit anti-mouse RELM**α** (Peprotech) and biotinylated goat anti-mouse Ym1 (R&D), followed by a 45min incubation with rabbit Alexa-Fluor 488 or AF594 Zenon labeling kit (Thermo) and Streptavidin PercP or BV421, BV785 or PeCy7.

For intracellular cytokine staining, cells were stimulated for 4h at 37°C with PMA (phorbol myristate acetate; 0.5µg/mL) and ionomycin (1µg/mL) and for 3h at 37°C with brefaldin A (10µg/mL; Biolegend). Cell surfaces were stained, and cells fixed as described above. All cells were permeabilized (Biolegend) then stained with antibodies for intracellular cytokines (**Supplementary Table 2**). Immune cells were identified according to the gating strategy in Supplementary figure 6. All samples were acquired with a FACSCanto II or 5 laser Fortessa with BD FACSDiva software and analysed with FlowJo software (versions 9 and 10; Treestar).

### RNA extraction and RT-qPCR

One right lung lobe was stored in RNAlater (Thermo) prior to homogenization in Trizol reagent (Thermo). RNA was prepared according to manufacturer’s instructions and stored at −70°C. Total RNA (0.5µg) was reverse transcribed using 50U Tetro reverse transcriptase (Bioline), 40mM dNTPs (Promega), 0.5µg primer for cDNA synthesis (Jena Bioscience) and recombinant RNasin inhibitor (Promega). The transcripts for genes of interest were measured by real-time qPCR with a Lightcycler 480 II system (Roche) and a Brilliant III SYBR Green Master mix (Agilent) including specific primer pairs (**Supplementary Table 3**). mRNA amplification was analysed by second derivative maximum algorithm (LightCycler 480 Sw 1.5; Roche) and expression of the gene of interest was normalised to a combination of housekeeping genes as specified in each figure.

### Histology and Immunostaining

The left lung lobe was fixed perfused with 10% neutral buffered formalin (Sigma) and was incubated overnight before being transferred to 70% ethanol. Lungs were processed and embedded in paraffin, then sectioned (5µm) and stained with Masson’s trichrome (MT), picosirius red or periodic acid schiff (PAS) stains using standard protocols. Images were captured with an 3D Histech Pannoramic P250 slide scanner or for Sirius red staining, collagen was imaged under brightfield and polarised light with an Olympus BX63 microscope. Analysis of Masson’s trichrome area and PAS+ area were analysed using a pixel classifier in QuPath (version 0.4.2). ROIs drawn around small to medium sized airways in the lung section were used limit measurements to the airways. Areas of staining were normalised to basement membrane length, and between 10-20 airways were analysed per mouse. Polarised images of sirius red staining was measured in airway ROIs limited areas to airways utilising the picrosirius red birefringence analyser script with Fiji (version 2.9.0/1.54b)^60^ (https://github.com/TCox-Lab/PicRed_Biref#readme). Briefly, a threshold was applied to picrosirius red birefringent signal to split each image into red-orange, yellow and green channels, corresponding to high, medium and low density/bundling of collagen fibres, respectively.

For immunostaining with antibodies or binding proteins, lung sections were deparaffinised and heat-mediated antigen retrieval performed using Tris (10mM) EDTA (1mM), Tween-20 (0.05%) buffer pH 9.0 (20min 95°C). Non-specific protein was blocked with 2% normal donkey serum in PBS containing 1% BSA and 0.05% Tween-20. If a biotin labelled antibody or binding protein was used, avidin biotin block (Biolegend) was performed prior to an overnight incubation at 4°C with primary antibodies and binding proteins (Supplementary Table 3). Sections were washed in PBS before incubation with secondary antibodies or labelled streptavidin (Supplementary Table 4) for 1hr at room temperature followed by mounting with DAPI containing fluoromount (Southern Biotech). Small to medium sized airways in lung sections were captured by an investigator blind to sample identity with an EVOS FL imaging system (Thermo). Images were analysed with Fiji (version 2.9.0/1.54b) as described previously^37^. Briefly, autofluorescence background was subtracted from each image based on pixel intensity from images where sections were stained with secondary only. ROIs were drawn around the airway, 50µm from the basement membrane, and positive staining area and intensity measured on images with an applied threshold. Positive area was normalised to basement membrane length and 5-15 airways were measured per mouse.

### Mass Spectrometry

Protein concentrations were quantified for Bronchoalveolar lavage samples and normalised accordingly. 1mg of protein was then incubated with 500mM IAM for 30min protected from light, followed by acidification with 12% aqueous phosphoric acid to a final concentration of 1.2%. Samples were then centrifuged at 10,000 RCF for 5min to pellet any debris and the clean supernatant was transferred to a new tube. 600µL of s-trap binding buffer was added to 100µL of sample and added in 200µL increments to the micro S-Trap column (ProtiFi, C02-micro) and spun at 2,000*g* for 30sec between each loading. Samples were then washed twice with s-trap loading buffer before addition of 2µg of Trypsin and incubated for 1h on a shaking 47°C incubator (Eppendorf, ThermoMix C). Samples were then removed from the column in a three-step elution with 40µL of TEAB (pH 8) followed by 40µL of 0.2% formic acid in H_2_O and finally 40µL of 50% acetylnitrile with 0.2% formic acid followed by lyophilisation using a speed-vac (Heto Cooling System) and storage at 4°C until clean up. Clean up and desalting of the samples was started by resuspending lyophilised peptides in 100µL of 5% acetylnitrile with 0.1% formic acid. Then in a 96-well 0.2µm PVDF filter plate (Corning, 3504) 100µL of a 10mg/mL settled Oligo R3 resin (Thermo Scientific, 1-1339-03) in aqueous 50% acetonitrile was added and washed with aqueous 50% acetonitrile and spun at 200*g* for 30sec to remove supernatant followed by one wash each with 100µL of 50% acetylnitrile and 0.1% formic acid. Samples were then added to the washed beads and mixed on a plate shaker (Eppendorf, ThermoMix C) for 5min at 200*g*. Samples were then washed three times with 0.1% formic acid with incubation for 2min at 200*g* each time and centrifugation at 200*g* for 30sec to remove supernatant. Samples were then eluted using 50% acetylnitrile and lyophilised prior to storage at 4°C before analysis. Sample peptides were evaluated by liquid chromatography coupled tandem mass spectrometry using an UltiMate 3000 Rapid Separation LC system (Dionex Corporation) coupled to a Q Exactive HF mass spectrometer (Thermo Fisher Scientific). Raw spectra were aligned using MAXQuant software v1.6.17.0 ^95^ with “matched between runs” being enabled. Raw data were then imported into R for differential analysis with MSqRob^96^ using the default pipeline. Heat maps were plotted using scaled log10 transformed LFQ counts.

### Nanostring

Extracted RNA were quantified using a Qubit™ RNA BR Assay Kit (Thermofisher) kit and subsequently diluted to 20ng/μL in RNase free water. Sample were run on a Nanostring nCounter® FLEX system using the Myeloid Innate Immunity v2 panel (XT-CSO-MMII2-12) as per manufacturer’s instructions. Raw data were loaded into nSolver version 4.0 using default settings and non-normalised counts were extracted and loaded into R for subsequent analyses were performed in R (version 4.2.0) using RStudio Version 2023.03.0 Build 386 – © 2009–2023 RStudio, Inc. Internal housekeeping as well as positive and negative control probes were used to ensure data integrity and set thresholds for minimum expression. Data were normalised using the edgeR package ^97^ and then differential expression was calculated using the limma package with the function voomwithqualityscores(). Figures were generated in R using ggplot and the complexheatmap package. Heat maps were plotted using scaled log10 transformed counts normalised to housekeeping references.

### Statistical analysis

Statistical analysis was preformed using JMP Pro 12.2.0 for Mac OS X (SAS Institute Inc.) or Graphpad Prism 8. Normal distribution of data was determined by optical examination of residuals, and each group was tested for unequal variances using Welch’s test. Differences between groups were determined by analysis of variance (ANOVA) followed by a Tukey-Kramer HSD multiple comparison test or unpaired two-tailed Student’s t-test as indicated in figure legends. In some data sets, data were log transformed to achieve normal distribution. Comparison with a *P*-value of <0.05 were considered statistically significant.

## Acknowledgements

The authors thank Stella Pearson and Brian Chan for their technical support, Peter Cook for advice on flow cytometry, and Hannah Tompkins for critically reading the manuscript. We also thank the Flow Cytometry, Histology and Biological Services core facilities at the University of Manchester.

