## Supplemental Tables and Figures for "Allergen-induced airway matrix remodelling in mice can be prevented or reversed by targeting chitinase-like proteins"

**Supplementary Table 1**: Primers and probe sequences used for ddPCR.

| Gene | Forward Primer | Reverse Primer | Probe | Label |
|---|---|---|---|---|
| *Chia* | CATGATCTGGGCCATTG | AGGGCTTTGTTCAAAGTAG | ACTGGCTCTTTCTGTGATCAGGGA | FAM |
| *Chil1* | TTCCTGCGTTCTTATGG | GATCAGGGTGGAGAAATAC | CCTGGCTCTACCCTCGCTTAAGAG | FAM |
| *Chil3* | GGGATGATATAGAGGTTTAGG | TATTGGCATCTGGTCTTG | ATGGGAAACTAGCAGTAGCCACCA | FAM |
| *Chil4* | GAATCTACTGATTCCCATCTC | CAGCCATTCATTAACTTACAC | TTCAGTAAGGAGGCACAGGAAGAG | FAM |
| *Tfrc* | CTGGTCAGCTCATTATTAAAC | GAACTGGTTCAGATCCTTC | ACCCATGACGTTGAATTGAACCTGG | HEX |

**Supplementary Table 2**: Antibodies and labelling reagents used in flow cytometry.

| Antigen | Label | Company | Clone | Isotype |
|---|---|---|---|---|
| CD4 | AF700 | Biolegend | GK1.5 | Rat IgG2b, **κ** |
| Ly6C | AF700 | Biolegend | HK1.4 | Rat IgG2c, **κ** |
| TCRb | AF780 | eBioscience | H57-597 | Armenian Hamster IgG |
| Strep | APC-eF780 | eBioscience | - | - |
| CD3 | APCCy7 | Biolegend | 145-2c11 | Armenian Hamster IgG |
| CD19 | APCCy7 | Biolegend | 6d5 | Rat IgG2a, **κ** |
| NK1.1 | APCCy7 | Biolegend | PK136 | Mouse IgG2a, **κ** |
| Ter119 | APCCy7 | Biolegend | TER-119 | Rat IgG2b, **κ** |
| MerTK | APC | Biolegend | 2B10C42 | Rat IgG2a, **κ** |
| IL-5 | APC | Biolegend | TRFK5 | Rat IgG1, **κ** |
| Ym1 | Biotin | R&D | Polyclonal | Goat IgG |
| B220 | Biotin | Biolegend | RA3-6B2 | Rat IgG2a k |
| CD11b | Biotin | Biolegend | M1/70 | Rat IgG2b k |
| CD11c | Biotin | Biolegend | N418 | Armenian Hamster IgG |
| Ter119 | Biotin | Biolegend | Ter-119 | Rat IgG2b k |
| NK1.1 | Biotin | Biolegend | PK136 | Mouse IgG2a K |
| Ly6G | Biotin | Biolegend | 1A8 | Rat IgG2a k |
| CD3 | Biotin | Biolegend | 17A2 | Rat IgG2b k |
| SigF | BV421 | BD Bioscience | E50-2440 | Mouse IgG2a K |
| CD11b | BV421 | Biolegend | M1/70 | Rat IgG2b k |
| CD11c | BV421 | Biolegend | N418 | Armenian Hamster IgG |
| Strep | 421 | Biolegend | - | - |
| IL-5 | BV421 | Biolegend | TRFK5 | Rat IgG1 k |
| IL-17A | BV510 | Biolegend | TC11-18H10.1 | Rat IgG1 k |
| MHCII | BV510 | Biolegend | M5/114.15.2 | Rat IgG2b k |
| CD90.2 | BV510 | Biolegend | 53-2.1 | Rat IgG2a k |
| CD11c | BV605 | Biolegend | N418 | Armenian Hamster IgG |
| CD3 | BV650 | Biolegend | 17A2 | Rat IgG2b k |

| | | | | |
|---|---|---|---|---|
| IFNg | BV711 | Biolegend | XMG1.2 | Rat IgG1 k |
| CD11b | BV711 | Biolegend | M1/70 | Rat IgG2b k |
| CD45 | BV785 | Biolegend | 30-F11 | Rat IgG2b k |
| Strep | BV785 | Biolegend | - | - |
| Zenon Rabbit IgG | F488 | Thermo | - | Rabbit IgG |
| IL-17A | F488 | Biolegend | TC11-18H10.1 | Rat IgG1 k |
| TCRgd | PE | Biolegend | GL3 | Armenian Hamster IgG |
| CD64 | PE | Biolegend | S18017D | Rat IgG2a k |
| IL-4 | PE/Dazzle594 | Biolegend | 11b11 | Rat IgG1 k |
| SigF | PE/CF594 | BD Bioscience | E50-2440 | Rat LOU |
| Zenon Rabbit IgG | AF594 | Thermo | - | Rabbit IgG |
| CD19 | Pecy5 | Biolegend | 6D5 | Rat IgG2a k |
| CD25 | PeCy5 | Biolegend | 3C7 | Rat IgG2b k |
| IL-13 | PeCy7 | eBioscience | eBio13A | Rat IgG1 k |
| Strep | Pecy7 | Biolegend | - | - |
| CD8 | PerCP/Cy5.5 | Biolegend | 53-6.7 | Rat IgG2a k |
| Ly6G | PerCP/Cy5.5 | Biolegend | 1A8 | Rat IgG2a k |
| RELMα | - | Peprotech | Polyclonal | Rabbit IgG |

**Supplementary Table 3**: Primer sequences for RT-qPCR.

| Gene | Forward Primer | Reverse Primer | Length (bp) |
|---|---|---|---|
| *Chil1* | CCAGCCAGGCAGAGAGAAAC | GCCACCTTTCCTGCTGACA | 57 |
| *Chil3* | TCTGGTGAAGGAAATGCGTAAA | GCAGCCTTGGAATGTCTTTCTC | 64 |
| *Chil4* | TCTGGTGCAGGAAATGCGTAAA | TCTGGTGCAGGAAATGCGTAAA | 64 |
| *Chia* | GTCTGGCTCTTCTGCTGAATGC | TCCATCAAACCCATACTGACGC | 377 |
| *Retnla* | TATGAACAGATGGGCCTCCT | GGCAGTTGCAAGTATCTCCAC | 107 |
| *Il4* | CCTGCTCTTCTTTCTCGAATGT | CACATCCATCTCCGTGCAT | 127 |
| *Il5* | ACATTGACCGCCAAAAAGAG | CACCATGGAGCAGCTCAG | 136 |
| *Lox* | CACTGCACACACACAGGGAT | TGTCCAAACACCAGGTACGG | 283 |
| *Rpl13a* | CATGAGGTCGGGTGGAAGTA | GCCTGTTTCCGTAACCTCAA | 116 |
| *Gapdh* | CATCACTGCCACCCAGAAGACTG | ATGCCAGTGAGCTTCCCGTTCAG | 153 |
| *Rn45s* | GTAACCCGTTGAACCCCATT | GTAACCCGTTGAACCCCATT | 151 |

**Supplementary Table 4**: Antibodies and labelling reagents used in immunofluorescence.

| Antigen | Label | Company | Clone | Species |
|---|---|---|---|---|
| Ym1 | Biotin | R&D | Polyclonal | Goat IgG |
| Ym2 | - | Cambridge Research Biochemicals | Polyclonal | Rabbit IgG |
| Collagen I | - | Cambridge Bioscience | Polyclonal | Goat IgG |
| Collagen III | - | Peprotech | Polyclonal | Rabbit IgG |
| Collagen IV alpha 1 | - | Novus | Polyclonal | Rabbit IgG |
| HABP | Biotin | Sigma | - | - |
| αSMA | - | Abcam | Polyclonal | Goat IgG |
| Vimentin | - | Abcam | Polyclonal | Rabbit IgG |
| Lox | - | Abcam | EPR21203 | Rabbit IgG |
| Anti-Goat IgG | NL637 | Bio-Techne | - | Donkey |
| Anti-Goat IgG | NL557 | Bio-Techne | - | Donkey |
| Anti-Rabbit IgG | NL637 | Bio-Techne | - | Donkey |
| Anti-Rabbit IgG | NL557 | Bio-Techne | - | Donkey |
| Streptavidin | NL637 | Bio-Techne | - | - |
| Streptavidin | NL557 | Bio-Techne | - | - |

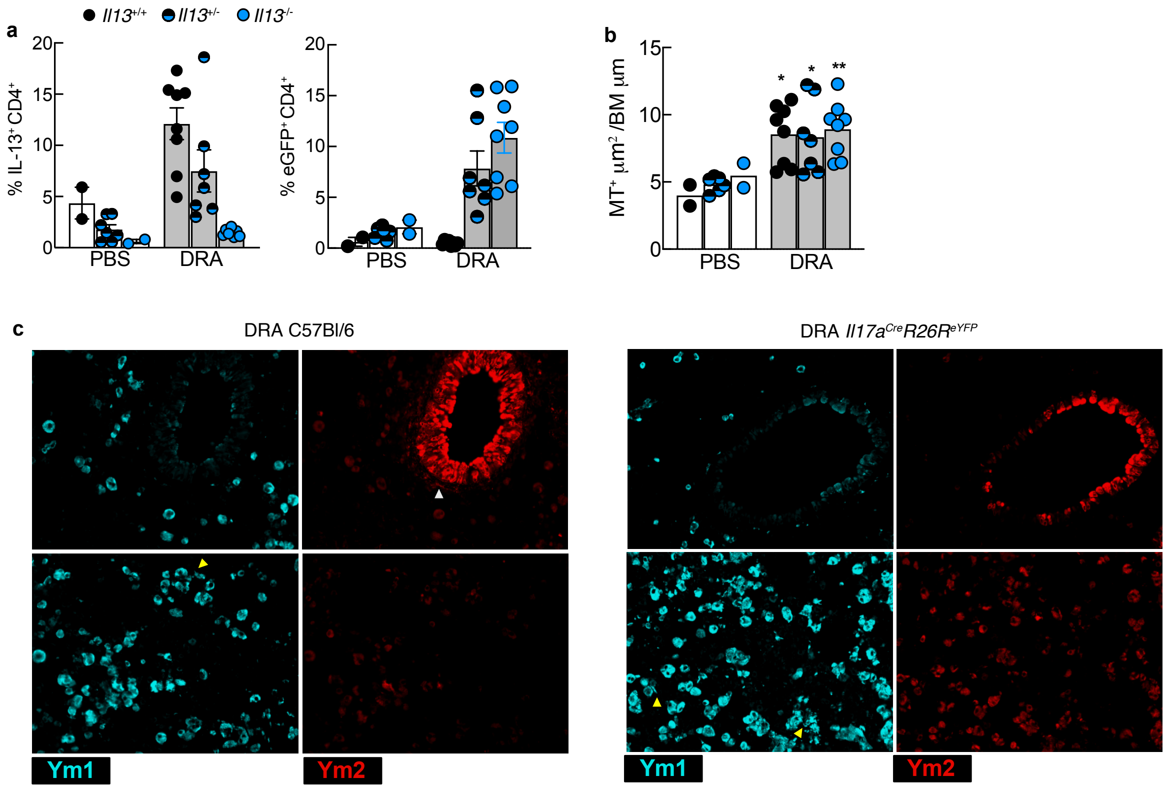


### Supplementary Figure 1

**Chitinase-like protein expression during IL-13 and Il-17a-independently allergic pathology**

**a**) *Il13*$^{eGFP}$ wild-type, heterozygote or homozygote mice were treated with PBS and DRA intranasally twice weekly for 8 weeks. Single cell suspensions from lungs were stimulated with PMA/ionomycin prior to staining cells for flow cytometry. Percentage of CD4$^{+}$ TCRβ$^{+}$ cells expressing IL-13 or eGFP signal in the lungs. **b**) Lungs from *Il13*$^{eGFP}$ littermate mice from a) were sectioned and stained with Masson's trichrome and total collagen accumulation around the airway was measured and normalised to basement membrane length. **c**) C57BL/6 or *Il17a*$^{Cre}$ *R26*$^{eYFP}$ mice were treated with PBS and DRA intranasally twice weekly for 8 weeks. Lung sections were stained with Ym1 and Ym2 to visual positive cells in the parenchyma and airways and representative images are shown. Yellow triangle shows Ym1$^{+}$ crystal material and white triangles show Ym2$^{+}$ cells located in areas where deposition of the ECM occurs in allergic mice. Datapoints from a, b show individual animals with bars representing mean ± s.e.m with n=2-8 female mice per group and data are from two combined experiments.

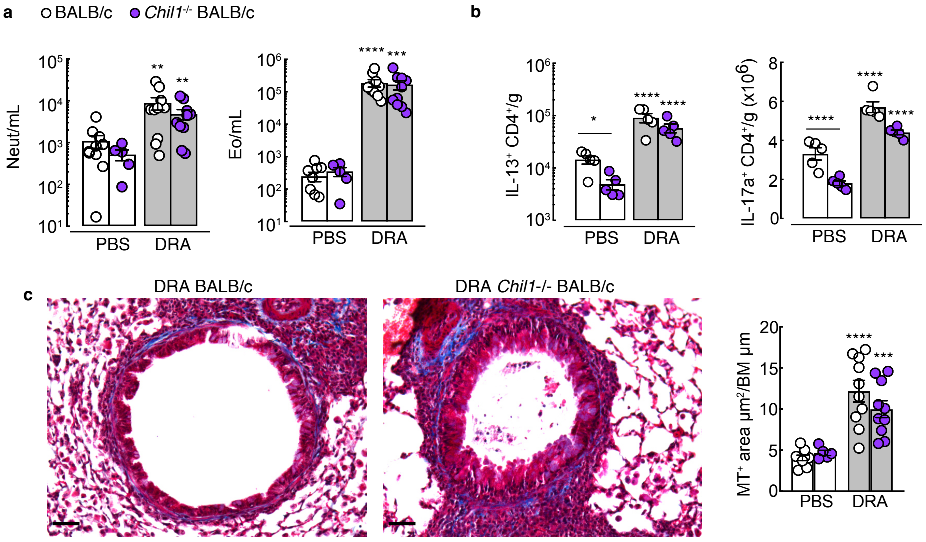


### Supplementary Figure 2

**Brp39 does not influence ECM remodeling in DRA mouse model of allergic pathology.**
**a**) *Chil1*$^{-/-}$ and BALB/c wild-type mice were treated with PBS and DRA intranasally twice weekly for 8 weeks. Cells were isolated from the BAL and numbers of neutrophils and eosinophils analysed by flow cytometry. **b**) Numbers of IL-13$^{+}$ and IL-17a$^{+}$ CD4+ TCRβ$^{+}$ cells per gram of lung were analysed by flow cytometry following PMA/ionomycin stimulation of single cell suspensions of lungs. **c**) Lung sections from mice in **a**) were stained for Masson's trichrome and total accumulation of collagen around the airways analysed. Datapoints show individual animals with bars representing mean ± s.e.m with n=5-10 female mice per group and data are from two combined experiments. Data were analysed by ANOVA with Tukey's multiple comparison test and significance level shown relative to PBS BALB/c mice as indicated on the graph. *$P$<0.05, ** $P$<0.01, *** $P$<0.001 **** $P$<0.0001 and NS, not significant.

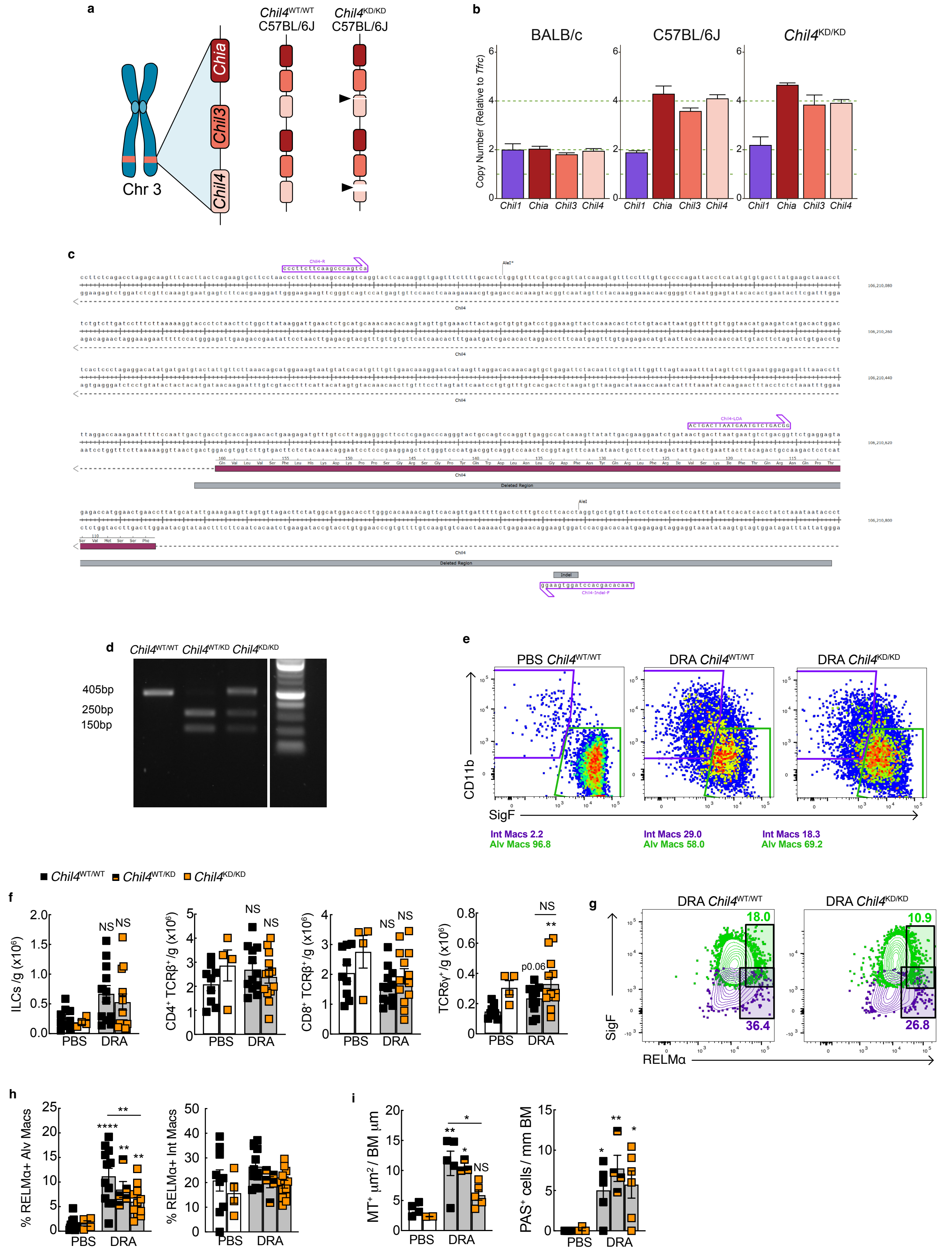

Supplementary Figure 3

**Development of *Chil4*$^{KD}$ mouse.**

**a**) Schematic diagram illustrating the location of *Chia*, *Chil3* and *Chil4* on chromosome 3 of the reference genome and the duplication of this region within C57BL/6J mice. Triangles in *Chil4*$^{KD/KD}$ mice show deletion of target exon 5 on one copy of *Chil4*, and deletion of a 6bp indel in the other *Chil4* gene copy. **b**) Digital droplet PCR analysis of *Chil1*, *Chia*, *Chil3* and *Chil4* copy number in BALB/c, C57BL/6J and *Chil4*$^{KD/KD}$ mice. Copy number of genes were measured relative to *Tfrc* known to have a copy number of 2. **c**) Partial map of the Chil4 gene with exon 5 highlighted in purple and deleted regions including the indel in grey. Primer sequences and AleI* restriction enzyme site used for genotyping are also shown in the diagram. **d**) Representative DNA gel showing genotyping results for Chil4 transgenic mice. **e**) Flow cytometry plots of MerTK$^{+}$ CD64$^{+}$ cells from the lungs of *Chil4*$^{KD}$ littermate mice treated with PBS or DRA for 8 weeks and stained with CD11b and SigF. Green boxes highlight alveolar macrophages whilst purple boxes highlight interstitial macrophages. **f**) Total numbers of ILCs, CD4$^{+}$ T cells, CD8$^{+}$ T cells and gamma delta T cells in the lungs of mice from d) as assessed by flow cytometry analysis. **g**) Representative flow plots and **h**) corresponding analysis of intracellular staining for RELMα within Alv.MØ and Int.MØ from mice as in **d**), analysed by flow cytometry with analysis. Plots are representative of n=12 mice per group from two combined experiments. Numbers within the box indicate percentage of cells positive for RELMα$^{+}$. Lung sections from mice in **d**) were stained for **h**) Masson's trichrome or PAS and total accumulation of collagen around the airways or numbers of PAS$^{+}$ cells analysed. Datapoints show individual animals with bars representing mean ± s.e.m with n=4-10 female mice per group and data are from two combined experiments.

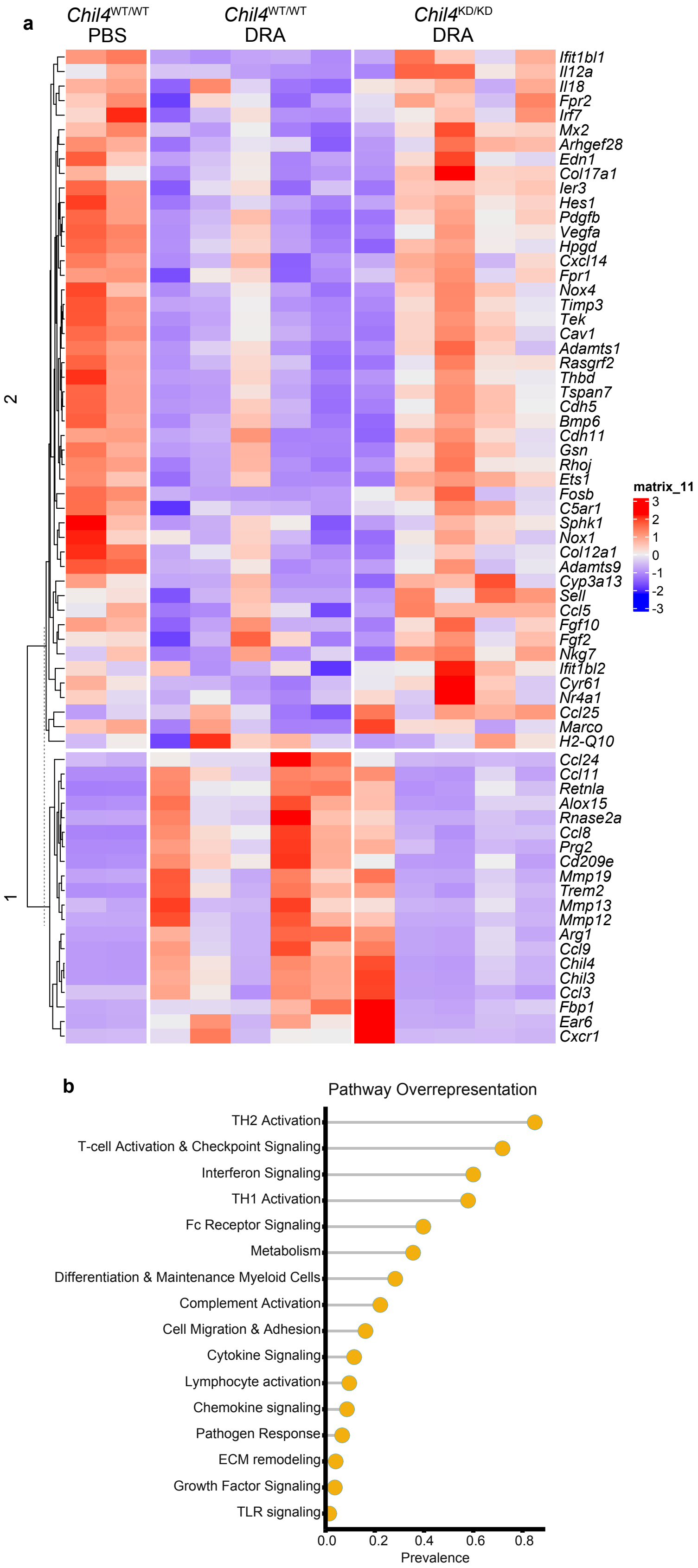


### Supplementary Figure 4

**Regulation of immune response genes in the lungs of *Chil4*[KD] mice.**

Whole lung RNAs from *Chil4*[KD] littermate mice administered PBS or DRA for 8 weeks were analysed with Nanostring Myeloid Panel version 2. **a**) Unsupervised, hierarchically clustered heatmap of genes that were significantly regulated in *Chil4*[KD/KD] DRA mice compared with *Chil4*[WT/WT] DRA mice. Corresponding genes from

*Chil4*$^{WT/WT}$ mice treated with PBS also shown. **b**) Analysis of significantly upregulated genes with pathway analysis.

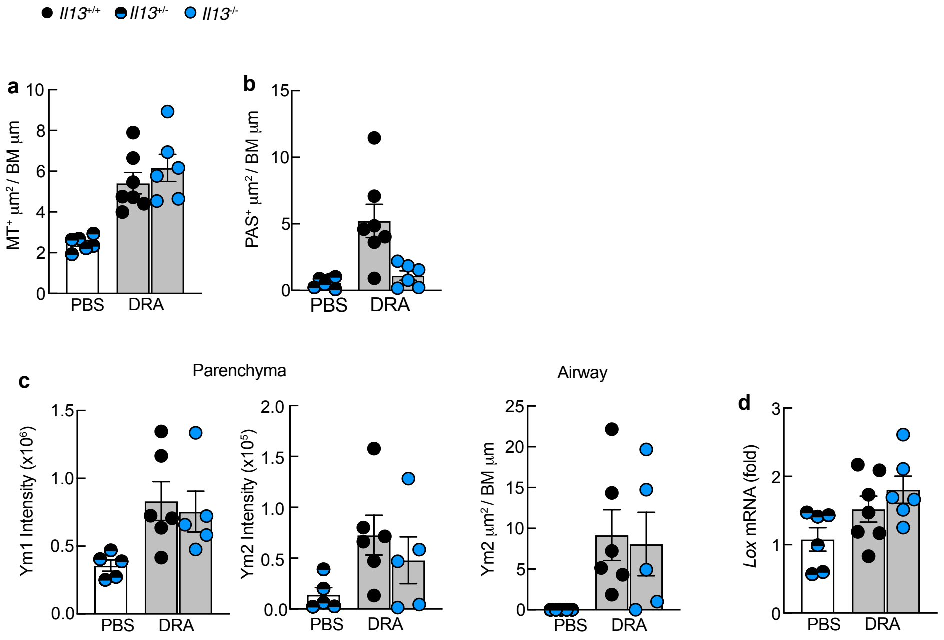


### Supplementary Figure 5

**Il-13 does not regulate pulmonary expression of lysyl oxidase.**

*Il13*$^{eGFP}$ wild-type, heterozygote or homozygote mice were treated with PBS or DRA intranasally twice weekly for 2 weeks. **a**) Lung sections were stained with Masson's trichrome and total collagen accumulation around the airways measured relative to basement membrane length. **b**) Numbers of PAS$^{+}$ airway epithelial cells normalised to basement membrane length in lung sections stained with PAS. **c**) Lung sections were stained with Ym1 and Ym2 and intensity of staining in the parenchyma or airways analysed. **d**) whole lung mRNA expression of Lox normalised to geometric mean of housekeeping genes *Gapdh*, *Rpl13a* and *Rn45s.*

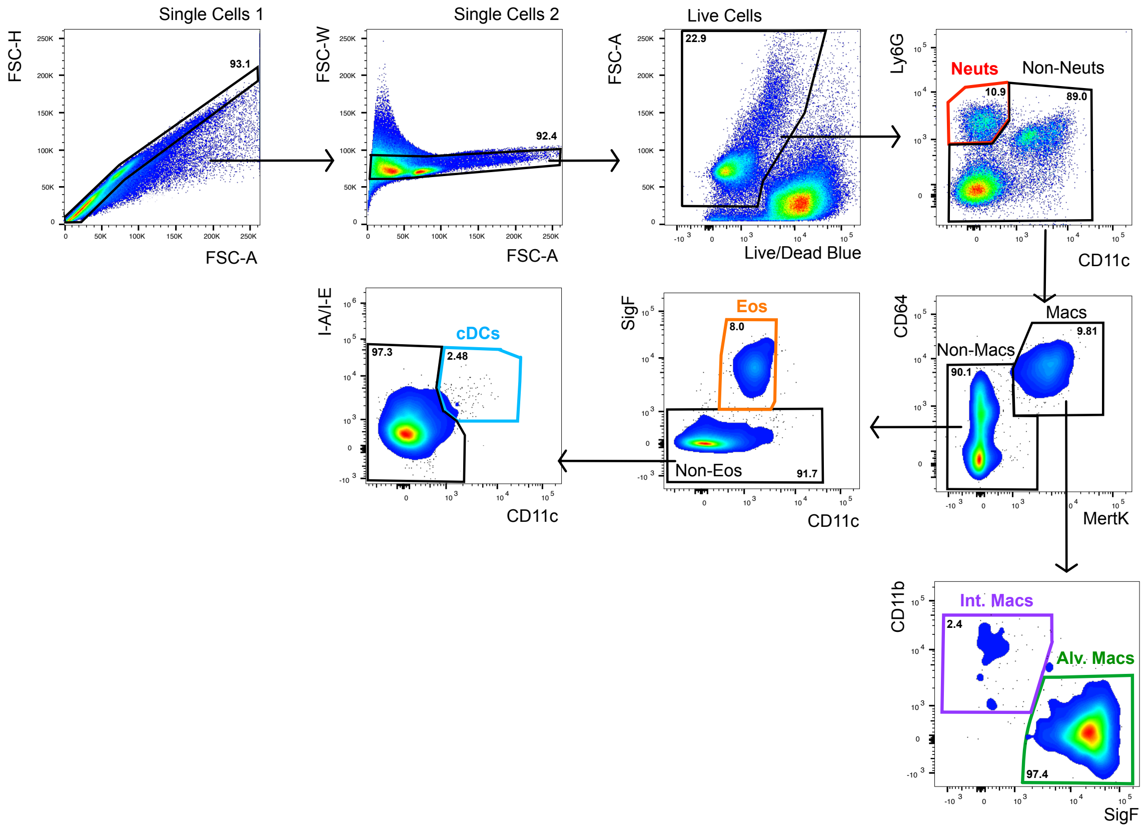


### Supplementary Figure 6

#### Gating strategy.

Flow cytometry was used to identify different immune cell populations in the lungs.
